# Distinct transcriptional states of draining lymph node T cells following influenza vaccination in individuals with underrepresented ancestries

**DOI:** 10.64898/2026.09.19.752896

**Authors:** Seungjoon Kim, Jacqueline H. Y. Siu, Terrence Chan, Charandeep Kaur, Chloe Hyun-Jung Lee, Wanyue Zhang, Jonas Mackerodt, Tamas Szommer, Kyla B. Dooley, Sofia Coelho, Aime Palomeras, Jamie Fowler, Brian D. Marsden, Teresa Lambe, Donald B. Palmer, Hashem Koohy, Nicholas M. Provine, Mark Coles, Calliope A. Dendrou, Katrina M. Pollock

## Abstract

The transcriptional and cell repertoire changes of lymph node (LN) T cells following influenza vaccination have been under-investigated across different ethnicities. Draining (dLN) and non-draining (ndLN) axillary LN cells from 17 participants with genotypically varied African or Asian ancestry (ISRCTN13657999) were collected by fine needle aspiration (FNA) before, 5 days and 42 days after adjuvanted influenza vaccination and analysed by single-cell RNA-sequencing (scRNA-seq). The early dLN transcriptional profile and cellular dynamics were distinct from the ndLN; BACH2 expressing T cells were highly induced, exhibiting stem-like and unique quiescence signatures. Proliferating T follicular helper cells (Tfh) were enriched in dLN and Th1-like Tfh predominated with highly induced transcriptional changes. The BACH2^+^ T cell state and Th1-like Tfh cells, transcriptionally distinct from germinal centre (GC) T cells, were co-localised with LAMP3^+^ dendritic cells in the T cell zone of publicly available LN datasets, a region that was also distinct from GC Tfh niches. Together, these findings indicate that the induction of a robust, proliferative, and transcriptionally diverse Tfh response, alongside a discrete subset of BACH2^+^ T cells, represents a marked feature of the early T cell zone immune response to adjuvanted influenza vaccine in the dLN.

## Introduction

Lymph nodes (LNs) are the critical organs that orchestrate the adaptive immune response, yet they remain largely understudied *in vivo* in humans during a functional immune response and have not been well described in adults of different ethnicities^1–3^. The global population is not fully represented by efforts to characterise immune cells at the single cell level, such as the Human Cell Atlas, and LN cells are also underrepresented, leaving two parallel critical knowledge gaps^4,5^. New personalised approaches to immunotherapy increasingly rely on an individual’s immune genotype and phenotype and there is a need for vaccines that protect all ancestral groups^6,7^. This knowledge gap becomes increasingly detrimental for groups who remain underrepresented in immunity research. At the same time, most LN vaccine research has focused on the germinal centre (GC) B cell response, with comparatively less know about the T cell response, particularly in the extrafollicular region of the LNs, which includes the areas where T cells are abundant in the paracortical T cell zone^8,9^. Given this region is important for the early recall response to infectious antigens and has a role in protection against cancer and the development of autoimmunity, there is a need to better understand the early processes that initiate T cell immune memory in the LN^10,11^.

There is a dearth of data on the relationship between the response to influenza vaccine and the ancestry of the recipient, with most studies focusing on vaccine uptake, rather than vaccine mechanism. Influenza vaccine uptake data can differ across ethnic groups depending on the setting^12,13^. Whilst there is a plausible relationship between several vaccine-induced immune signaling pathways, including HLA type and ancestry, few have been rigorously investigated in vaccine studies. Reporting of genetic ancestry and self-declared ethnicity may be conflated with other social parameters including identity, leading to weaknesses in the way data are collected. Thus, a lack of specifically designed studies mean that African and Asian populations are underrepresented in immunology research.

Traditionally, our understanding of functional T cell differentiation has relied heavily on animal models, in which secondary lymphoid organs (SLOs), such as lymph nodes, provide the critical microenvironment for lymphocyte priming and activation^14^. T cells are enriched in the paracortex, where they undergo priming by antigenic stimulation by antigen presenting cells (APCs). After priming, naive T cells undergo clonal expansion and assume phenotypically and functionally distinct differentiation states with different capacities for memory and effector functions. However, T cells can also undergo quiescence, anergy, senescence, and exhaustion^15–17^. The differentiation of effector and memory T cells is governed by transcription factors including the T-bet/EOMES and BLIMP1/BCL-6 axes. T-bet and BLIMP1 are increased in effector cells while EOMES and BCL-6 are increased in memory cells acting as molecular rheostats, which coordinate CD8+ T cell fate^18^.

To develop new immune cell therapies and infectious disease or cancer vaccines for all ancestries, requires a better understanding of LN function at the single cell level. Immunisation with a licensed vaccine provides an *in vivo* perturbation that can be directly measured by sampling the draining (dLN) and non-draining lymph node (ndLN) using ultrasound (US) guided fine needle aspiration (FNA) in the axillae. Using this approach, we previously showed that the LN response to adjuvanted influenza vaccine is anatomically and temporally regulated within the dLN compared with the ndLN in an ancestrally diverse cohort of young adults. Specifically, T follicular helper cells (Tfh) were induced within five days within the dLN, and the gene expression programmes (GEPs) of induced LN cells formed transcriptional hubs that were not present in the ndLN^19^.

To contribute to addressing the under representation of different ancestries in immunology and vaccine research, and to understand cellular and transcriptional dLN and ndLN function in this context, we used seasonal influenza vaccine as a probe to stimulate a functional T-cell immune response in the human LN. Here we describe findings from the second cohort of a LN study of young healthy adults with self-declared African and Asian ancestry conducted in West London, UK (ISRCTN13657999) who were immunised with adjuvanted quadrivalent seasonal influenza vaccine (aQIV) in the winter season of 2023-2024. We demonstrate that induction of a distinct Th1-like subset of Tfh and upregulation of the transcription factor *BACH2* are dominant features of the dLN, but not the ndLN T cell response to immunisation. These T cell states were transcriptionally and spatially distinct from the GC response and co-located, extrafollicularly in the T cell zone of the LN in publicly available data. Our results raise the possibility that Th1-like Tfh and BACH2 responses form part of the early T cell immune response, and could originate within the extrafollicular region of the dLN in the first few days following aQIV immunisation in young African and Asian adults. These data form a publicly available resource which contributes to addressing inequality in immunology and vaccine research.

## Results

### Genotyping and LN cellular composition of rarely studied ancestrally diverse participants responding to aQIV

LN cells (LNCs) were obtained by FNA before immunisation (V2), at day 5 (V4) and at week 6-10 (V6) from 17 participants who completed the study protocol (female n = 15, male n = 2) (Figure 1A and Table S1). LNCs were analysed by single-cell RNA-sequencing (scRNA-seq) and cellular indexing of transcriptomes and epitopes by sequencing (CITE-seq). After batch correction and QC filtering, a total of 521,354 high-quality immune cells were retained alongside transcriptomes, cell-surface protein and T-cell receptor (TCR) repertoire (Figure 1B). Genetic ancestry inference was conducted against a reference dataset of genotypes (1000 Genomes Project^20^ and Human Genome Diversity Project^21^) applying ADMIXTURE^22^ to estimate individual ancestry proportions. The ancestry proportions were estimated for seven ancestral super populations (Figure 1C and S1A). By the largest individual ancestry proportion, participants grouped into four super populations (African, East Asian, Middle Eastern and Central/South Asian). This closely recapitulated the participant’s self-reported ancestry (Table S2) and underpins the diversity which is an important feature of the present work^1,4,6^.

**Figure 1.**
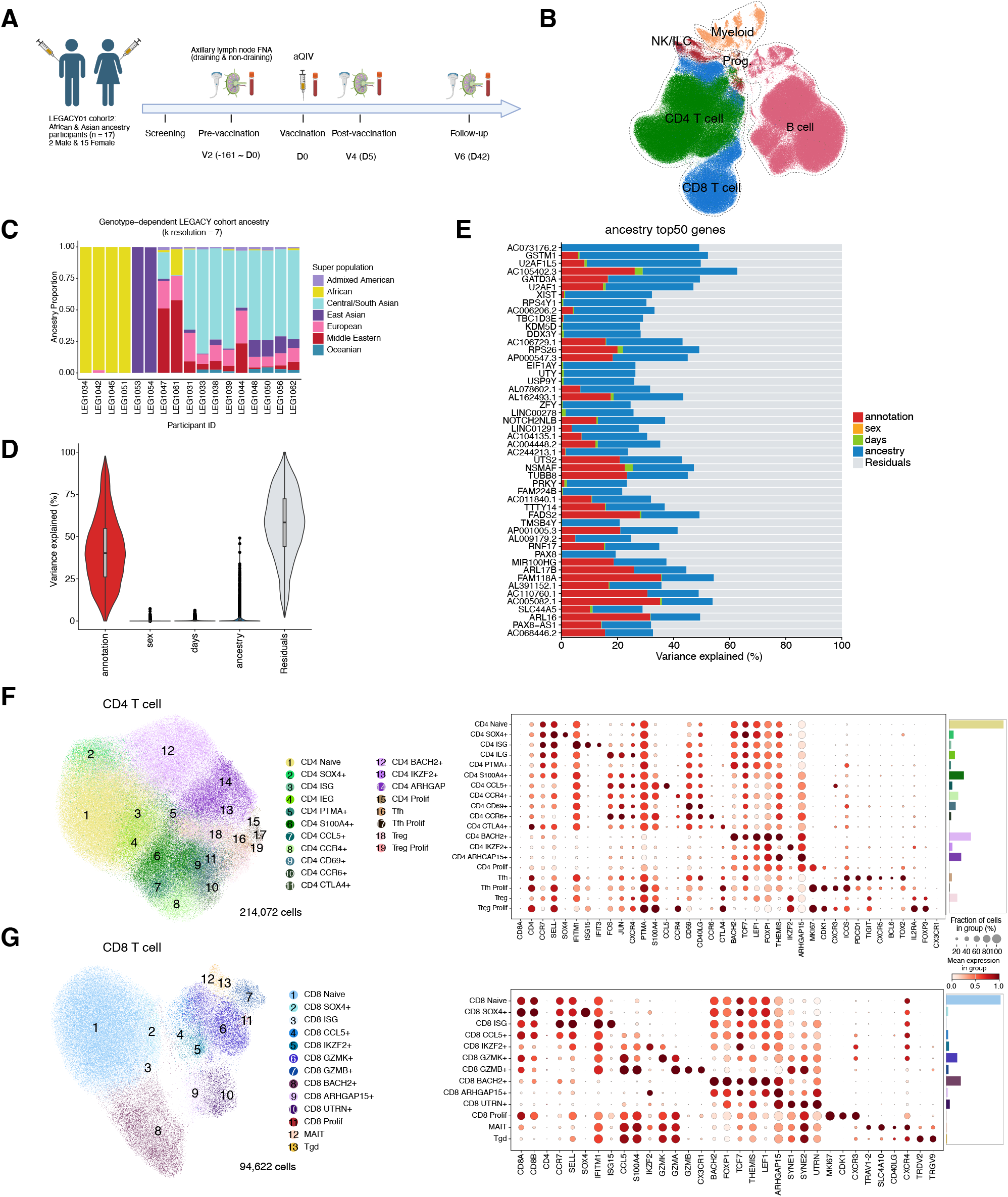
LEGACY study design and characterisation of LN FNA cell populations. (A) Longitudinal sample collection pre- (V2) and post-immunisation (V4 and V6). (B) UMAP representation of the broad immune cell compartments. (C) Stacked bar plot of global ancestry proportions for each individual participant by genotype. (D) Distribution of the percentage of variance assigned to each covariate across broad immune cells. (E) Percentage of variance assigned to each variable in the variance partition analysis. (F) UMAP of CD4^+^ T cells and (G) CD8^+^ T cells with key RNA marker genes (right) of each cell type.

Participants had human leucocyte antigen (HLA) genotype diversity, *HLA-A* (n = 7), *HLA-B* (n = 7), *HLA-C* (n = 2) and *HLA-DRB* (n = 5) alleles that were not in the top 10 alleles found in the public Europe allele frequency database^23^ (Table S2). Transcriptional variation across pseudobulk transcriptome of each gene among contributing sources including age, sex, days (pre and post immunisation) and ancestry was assessed. Most of the variance was explained by residuals or hidden covariates not identified within the study (Figure 1D). Ancestry explained an average of 0.92% of the variation and 37 genes accounted for more than 20% of the variation in the model. The top 50 example genes are shown (Figure 1E and S1B). For example, *GSTM1*, *AP000547.3* and *AL162493.1* exhibited higher expression levels in LNCs from those with African ancestry while *AC106729.1*, *LINC01291* and *AC104135.1* exhibited higher expression levels in East Asian ancestry than others (Figure S1B).

To explore T and innate lymphoid cell (ILC) dynamics in LNs, we isolated high quality T/ILC subsets from the dataset. Subsequent sub-clustering identified three main lineage compartments, including CD4^+^ and CD8^+^ T cells, and both natural killer (NK)/ILC populations. After annotation based on marker genes and label transfer from the cohort 1 study^19^, CD4^+^ T cells were sub-clustered into 19 different cell states (n = 214,072 cells), and CD8^+^ T cells were sub-clustered into 13 different cell states (n = 94,622 cells) (Figure 1F-G). Amongst both CD4^+^ and CD8^+^ T cells, in additional to classical cell states, there were cell types that expressed high levels of the transcription factor (TF) *BACH2* and the gene *ARHGAP15*, respectively. These cell type clusters were distinguished by low ribosomal gene expression and low expression of TRA/TRB locus genes compared with other T cell states (Figure 1F-G and Figure S2A-C). BACH2^+^ T cell states had several shared characteristics, including the expression of quiescent or stem-like genes including *TCF7*, *LEF1* and *FOXP1* and this may be characteristic of a quiescent, less differentiated cell state^24–26^. Protein expression of the chemokine receptors CXCR4, CXCR3 and CX3CR1 was high but gene expression of *CXCR4*, *CXCR3* and *CX3CR1* was low (Figure S2D-E). ARHGAP15^+^ T cell states had similar characteristics to BACH2^+^ T cell states except for the expression of stem-like genes (Figure 1F-G and Figure S2A-E).

NK/ILC cells were sub-clustered into 12 different cell states (n = 8,677 cells) and ARHGAP15^+^ NK and ILC cell-types also expressed *BACH2* and stem-like genes, suggesting poor separation of the two populations reflecting low numbers of these cell types (Figure S3A-C). The NK SYNE1^+^ state was also identified as the NK1B population from an NK atlas^27^, suggesting consistent sub-clustering between the two datasets (Figure S3A-B). The NK subsets exhibited distinct gene and protein expression profiles; while NK CD56^hi^ showed high expression of CD56 protein, NK ARHGAP15^+^ cell state expressed high levels of *NCAM1* (CD56 gene) despite exhibiting lower CD56 protein expression (Figure S3C).

To investigate the existence of the BACH2^+^ T cell state upon immunisation, we integrated and compared our data with published LN scRNA-seq data derived from a non-adjuvanted influenza vaccination study^28^. Cluster 10 from the earlier published study showed expression of *BACH2*, *ARHGAP15*, *LEF1*, *FOXP1* and *RUNX1,* and low ribosomal and TRA/TRB locus gene expression (Figure S4A-D). This cluster only existed at the early time points D12 and 26 (Figure S4E). Both cosine similarity and pearson distance demonstrated that the BACH2^+^ cell states we annotated were most like cluster 10 from the reference data (Figure S4F).

Transcriptional variation in each of the CD4^+^ and CD8^+^ T cell states by ancestry was assessed. Based on the top 100 ancestry variation genes, *PTGDR* was lowly expressed from those with East Asian ancestry and decreased upon vaccination in LNs CD4^+^ T cells, while *TNFSF8* was lowly expressed from those with African ancestry and increased upon vaccination in CD8^+^ T cells (Figure S5A-B). Further statistical characterisation of these differences was not possible due to the sample size. These findings indicated the unique genotyping and transcriptional characteristics of the LNCs were more heavily influenced by immunisation than ancestral diversity, with the BACH2^+^ T cell state emerging with a distinctive quiescent-like transcriptomic profile.

### Naive and BACH2^+^ T cell states were relatively enriched and Tfh was the most activated cell state in the dLN after vaccination

To evaluate how immunisation influences cell states in the dLN and the ndLN, cell type differential abundance (DA) and differentially expressed genes (DEGs) were performed. DA was analysed at the V2 (pre-vaccination) and V4 (early post-vaccination) timepoints using model-adjusted odds ratios through Bayesian hierarchical beta-binomial models. At V4, we observed a relative increase in abundance of proliferating Tfh, CD4^+^ and CD8^+^ naive T cells and BACH2^+^ T cells, alongside a trend towards an increased in Tfh after immunisation (Figure 2A). Conversely, ndLN exhibited relatively higher frequencies of CD4^+^ immediate early gene (IEG), ARHGAP15^+^ T cells, MAIT and NK cells as well as Treg and differentiated subsets such as CD4^+^ CCR4^+^ and CCR6^+^ T cells and CD8^+^ GZMK^+^ and GZMB^+^ T cells (Figure 2A). Model robustness was confirmed by repeating the model excluding the two participants who underwent the first post immunisation FNA later than 5 days, showing no overall differences (Figure S6A). The cell state differences were also seen in an analysis of cellular density along the post-vaccination days (Figure 2B, S6B-C).

**Figure 2.**
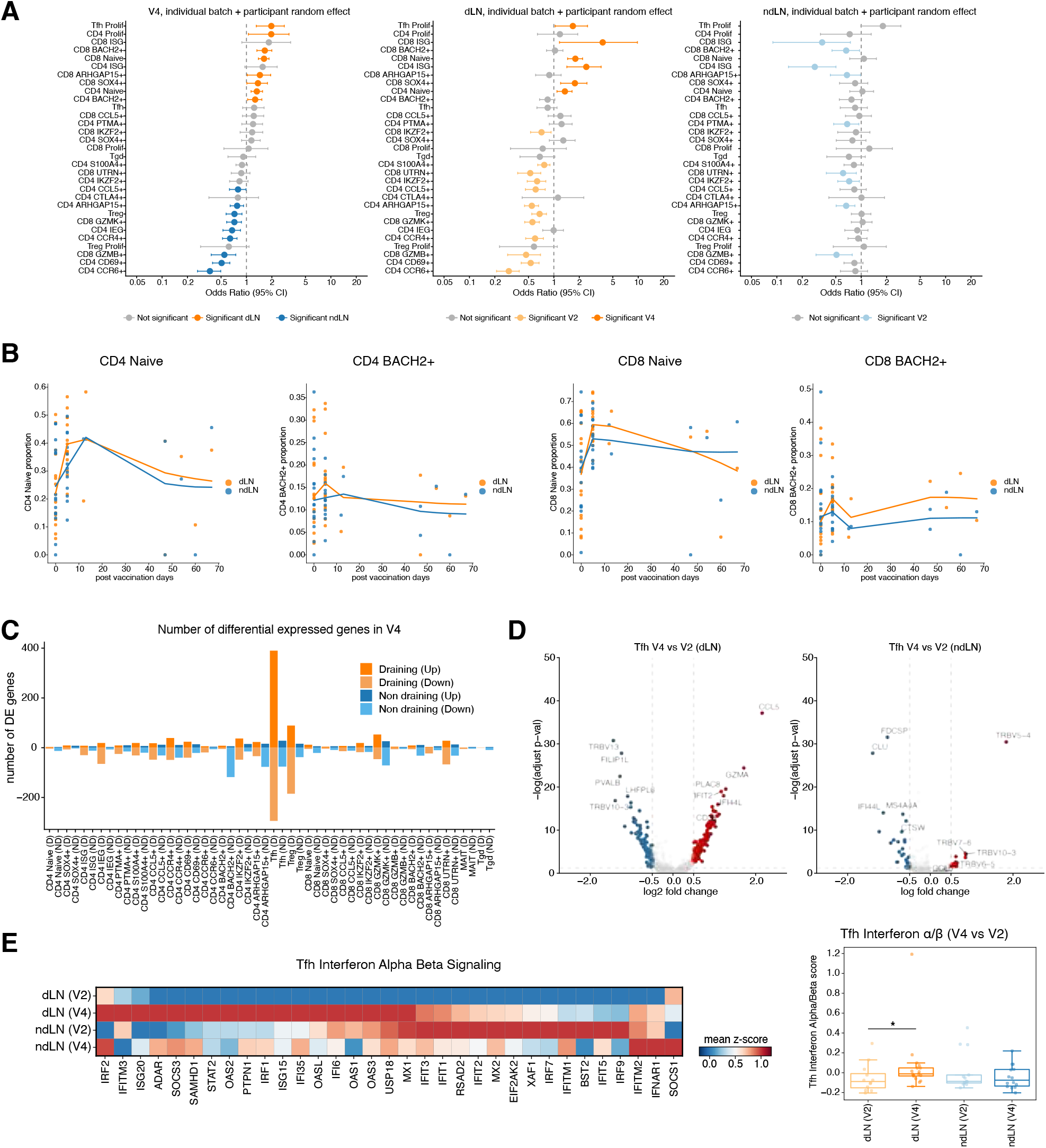
Proportion and transcriptional changes of LN FNA cell populations upon immunisation. (A) Differential abundance of dLN vs ndLN (left), V4 vs V2 at dLN (middle) and V4 vs V2 at ndLN (right) by beta-binomial model represented as odds ratios (OR). Non-grey dots indicate significant differences. Error bars show the 95% credible interval. (B) Cell proportion density along the post-vaccination days (left: CD4^+^ and right CD8^+^). (C) Bar plot showing number of DE genes (logFC > ± 0.5 and FDR < 0.05) per cell type. Positive is number of up-regulated while negative is number of down-regulated. (D) Volcano plots showing differentially regulated genes in Tfh across D4 vs D2 in dLN (left) and same in ndLN (right) (logFC > ± 0.5 and FDR < 0.05, blue: downregulated in V4; red: upregulated in V4). (E) Heatmap illustrating mean expression z-score for the genes from interferon alpha/beta signaling across Tfh cells grouped by different time points (left) and barplot showing interferon alpha/beta signaling score per each donor in different time points. Data were analysed using Wilcoxon rank-sum test and depicted as *p<0.05.

In DEGs analysis, Tfh within the dLN had the highest number of DEGs compared to other cell types, with a heightened transcriptional activation in the dLN relative to the ndLN at V4 post immunisation (Figure 2C). For example, *CD38* and multiple ISGs were activated in dLN Tfh but not in the ndLN Tfh. Also, *CCL5* was up-regulated across both anatomical sites at V4, with a higher log-fold change within the dLN (Figure 2D). Although increasing cell counts showed modest increase in number of DEGs (36%), cell type differences account for more variance (up: 136%, down: 58%) (Figure S7A and Table S3). To gain functional insight for each cell type, gene set enrichment analysis (GSEA) leveraging the Reactome gene sets^29^ was performed (Figure S7B). Interferon signals were up regulated in the dLN compared to ndLN, including in Naive cell types, CD4^+^ BACH2^+^ T cells and Tfh. Specifically, Tfh Interferon *a*/β signaling was significantly increased in the dLN, not in the ndLN (Figure 2E). Highly localised Defensin pathways were selectively down-regulated in a small number of CD4^+^ T cell subsets, but uniquely up-regulated within Tfh cells (Figure S7B). In contrast to these gene regulation dynamics, MAIT and γδT (Tgd) cells showed no distinctive pathway alterations between the dLN and ndLN at V4 (Figure S7B). Overall, upon aQIV immunisation, proliferating Tfh, naive and the BACH2^+^ T cell states were proportionally increased during the early response in the dLN. Amongst Tfh in total, there was a large change in the number of differentially expresses genes, especially interferon-related genes in the dLN.

### Delineation of Tfh subsets enriched in dLN upon immunisation

To comprehensively characterise Tfh, seven subsets were sub-clustered and their identities assigned based on the published literature of marker genes^28,30^ (Figure 3A). High expression of Tfh conventional markers, including *PDCD1*, *ICOS*, *TOX*, and *TOX2* compared to the rest of CD4^+^ T cells, confirmed their Tfh lineage^31^. Among the seven subsets, Th1-like Tfh cells were identified based on their expression of Th1 marker genes, including *TBX21*, *CCR5*, *CXCR3*, *CXCR6* and *IFNG*^30^. GC Tfh expressed high *PDCD1*, *ICOS*, *CXCR5* and *IL21*, and Tfh BACH2^+^ highly expressed *BACH2* and *FOXP1* relative to all other subsets. At the protein level, Th1-like Tfh cells expressed CCR5 and CXCR3 and ICOS and were transcriptionally distinct from GC Tfh (Figure 3A and S8A). To confirm these sub-clustering results, transcriptomic similarity analysis was performed comparing our annotations with published Tfh subsets from independent dLN scRNA-seq datasets^28^ (Figure S8B-C). The Th1-like Tfh cells were most similar to cell states annotated as Tfh cycling from the public data (Figure S8D-E), which expressed *CXCR3*, but lacked *CCR5* and *CXCR6* expression (Figure S8C). The cell states identified were thus concordant across studies.

**Figure 3.**
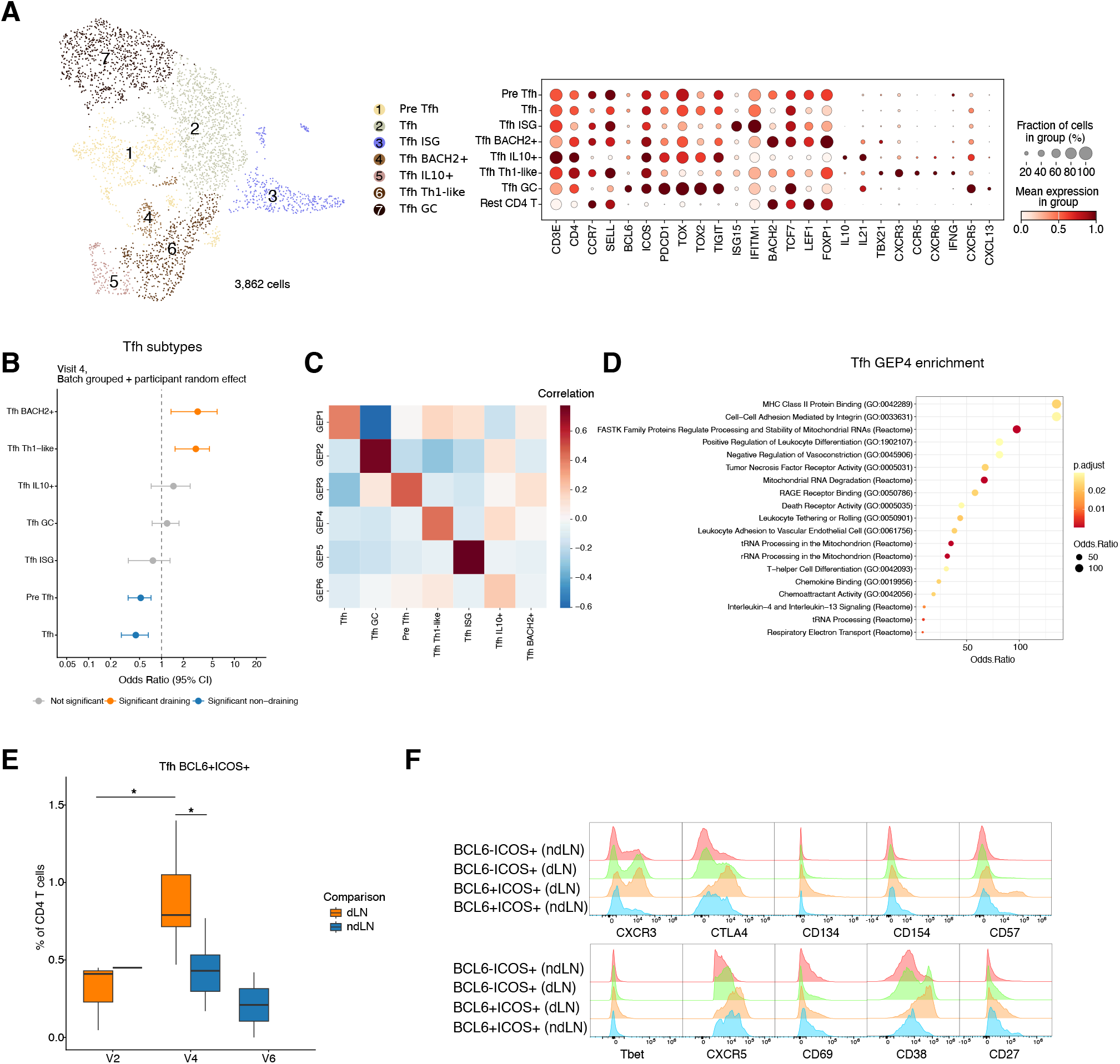
Distinctive Tfh subsets of human LN. (A) UMAP projection of Tfh subset annotations and dot plot of selected Tfh cell subset-specific markers (right) (B) Beta-binomial model was used to test abundance differences across cell compartments after influenza vaccination. Comparison of vaccination between V2 and V4 samples from dLN and ndLN as well as V4 comparison of dLN and ndLN (purple) represented as odds ratios (OR) are shown. Non-grey dots indicate significant differences (adjusted p-value < 0.05) after Benjamini-Hochberg correction. Error bars show the 95% confidence interval. (C) Identification of GEPs obtained using cNMF from Tfh cells. The color indicates a pearson correlation between each cell type. (D) Dot plot showing GO analysis of top 100 genes in GEP4. The dot size illustrates gene ratio, and the color denotes adjusted p-value value. (E) Barplot showing different proportions of BCL6+ICOS+ T cells per each donor in different time points. Data were analysed using Wilcoxon rank-sum test and depicted as *p<0.05. (F) Representative flow cytometry histograms showing the expression of the indicated markers on manually gated cell populations from concatenated samples.

Proliferating Tfh cells were more abundant in the dLN at V4, whilst pooled Tfh subsets did not reach statistical significance (Figure 2A). Repeating differential abundance analysis amongst Tfh subsets, Tfh BACH2^+^ and Th1-like Tfh subsets were relatively increased in the dLN, whilst pre-Tfh and other subsets decreased, indicating a change in Tfh subset composition with immunisation (Figure 3B). To understand phenotypic characteristics of Tfh subsets, we applied consensus non-negative matrix factorisation (cNMF) to Tfh subsets and identified 6 GEPs, amongst which GEP4 was highly correlated with the Th1-like Tfh cell state (Figure 3C). Based on its top gene feature profiles and the enriched pathways, GEP4 was involved in T cell antigen recognition and differentiation pathways and adhesion or chemokine binding pathways, consistent with the high expression of *CXCR3* (Figure 3D).

These findings were tested at the protein level by spectral flow cytometry of the same LNC samples where cell abundance allowed. ICOS⁺BCL6⁺ Tfh cells increased following immunisation, with higher frequencies in the dLN than in the ndLN (Figure 3E and S9A). FlowSOM clustering identified three Tfh clusters enriched in the dLN compared with the ndLN: C14 (CXCR3⁺, PD1^+^), C22 (CXCR3⁻, PD1^hi^), and C25 (CXCR3^hi^, CD38^+^), consistent with the enrichment of Th1-like Tfh cells observed by transcriptomic analysis (Figure S9B-C). ICOS⁺BCL6⁺ Tfh cells had higher expression of CTLA-4, CD134, CD57, CXCR5 and CD38 in the dLN compared with the ndLN, supporting a more activated Tfh phenotype in the dLN (Figure 3F and S9D). Taken together, gene expression and surface protein analyses of the same samples demonstrated the localised expansion of a CXCR3^+^ Tfh subset in the dLN upon immunisation.

### A distinctive BACH2^+^ T cell gene programme and TF regulon activity induced by immunisation

To characterise the observed T cell states in LNs, TF regulon activity and GEPs inference were performed. Based on pre-defined TF-target gene relationships, there was a clear distinction in regulatory profiles between classical T-cell lineages and putative quiescence lineages^16^ (Figure 4A). Regulons associated with stemness programs, including *SOX9*, *SOX4*, *TCF7*, *LEF1* and *FOXP1* were activated in these lineages^32^. SOX4 and SOX9 regulons were also activated in CD4^+^ SOX4^+^ and PTMA^+^ T cell states. CD4^+^ ARHGAP15^+^ T cells had high *ETV7* regulon activity which targets *MGAT5* known for terminal exhaustion^33,34^(Figure 4A). Tfh cells had activation of TBX21 and BATF regulons, as might be expected for Th1-like Tfh (Figure 4A). The dominant regulatory programs, defining each T cell state, were therefore as expected for classical cell types, and distinctive for BACH2^+^ and ARHGAP15^+^ T cell states (Figure S10A). Given *BACH2* and *ARHGAP15* expressing T cells shared similar TF profiles, we further investigated DEGs and pathways between the two states (Figure S10B). CD4^+^ BACH2^+^ T cells had high gene expression related to translation, viral infection and NOTCH signaling, while CD4^+^ ARHGAP15^+^ showed upregulation of pathways related to apoptosis, TNF signaling and cell migration mediated by Rho signaling (Figure S10C-D).

**Figure 4.**
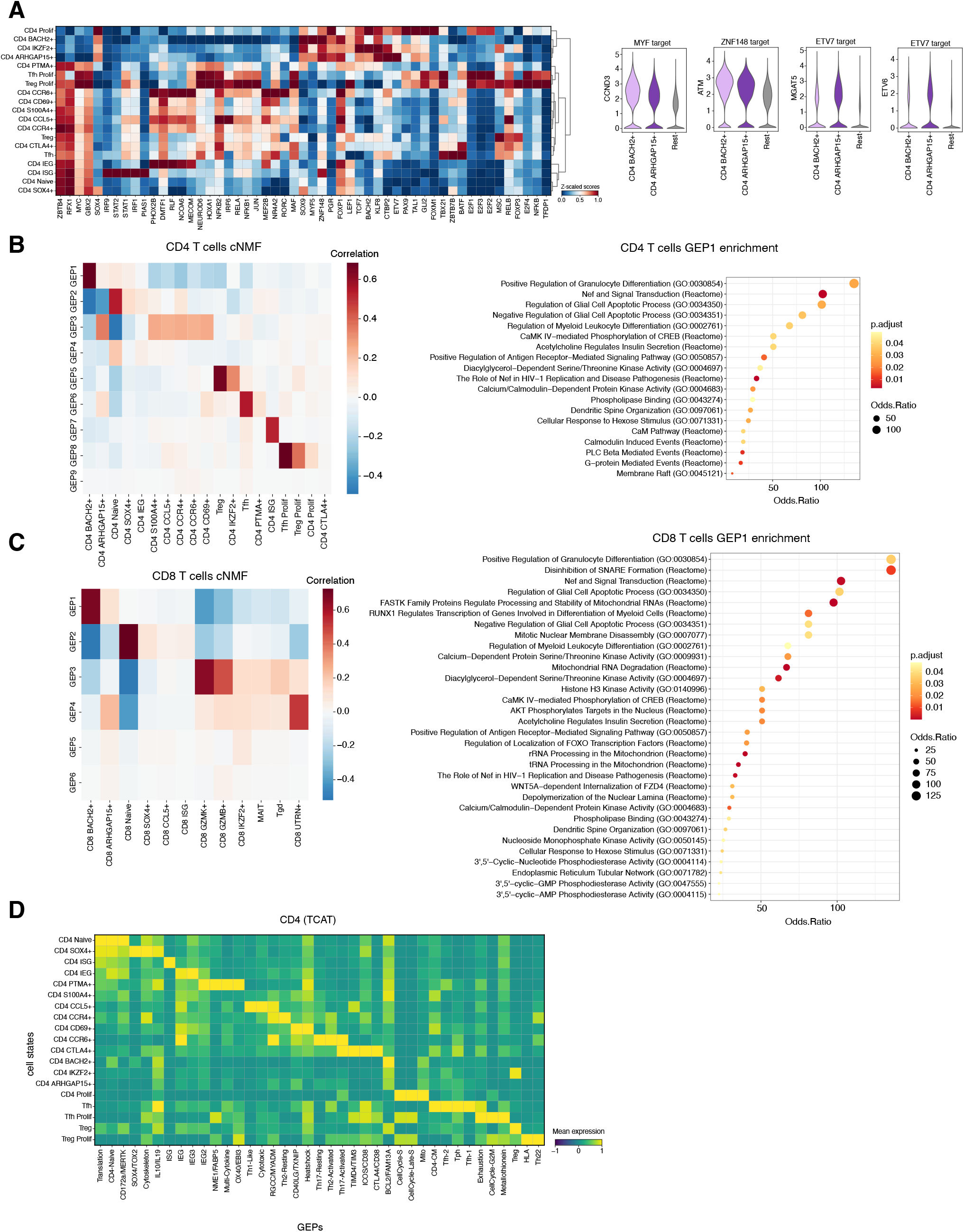
Transcriptional characterisations of LN FNA cell populations. (A) Heatmap illustrating the activity of TFs on each cell type from CD4 (left) and violin plots expressing representative target genes of TF (right). (B) Identification of transcriptional programs from cNMF on CD4^+^ and (C) CD8^+^ T cells. The color indicates a pearson correlation between each cell state. The dot plot (right) shows GO analysis of top 100 genes in GEP1 in each CD4^+^ and CD8^+^ T cell. From GO analysis, the dot size illustrates gene ratio, and the color denotes adjusted p-value. (D) Heatmap illustrating the GEPs from TCAT on each cell state from CD4^+^ T cell. The color indicates mean z-score per state.

Using cNMF, nine and six discrete GEPs were identified in CD4^+^ and CD8^+^ T cells (Figure 4B-C), respectively. From factor analysis in each lineage, GEP1 was highly correlated with the BACH2^+^ cell state. The top driving genes were extracted from GEP1 for functional enrichment analysis. GEP1 pathways were involved in granulocyte differentiation, glial cell apoptotic processes, insulin secretion, and serine/threonine or calcium/calmodulin-dependent protein kinase activities. HIV-1 related Nef-mediated pathways^35^ were significantly enriched, highlighting a conserved transcriptional state previously described in viral infection rather than immunisation (Figure 4B-C).

T-cell annotations in our dataset were benchmarked against public GEPs from the TCAT database^36^ and a pan-cancer T-cell atlas^37^. Most public GEPs aligned well with our annotations (Figure 4D). Notably, both CD4^+^ and CD8^+^ BACH2^+^ T cells were enriched for the BCL2/FAM13A program, which is an anti-apoptotic pathway that prevents mitochondrial membrane permeabilisation^38^ (Figure 4D and S11A-B). The pan-cancer atlas further revealed high expression of an anti-apoptotic signature within CD4^+^ IKZF2^+^ and CD8^+^ BACH2^+^ T cells (Figure S11C). Anergy gene sets were enriched in both CD8^+^ ARHGAP15^+^and UTRN^+^ cells, suggesting functionally inactivated phenotypes (Figure S11C).

Comparative analysis revealed similarities and differences in GEPs between the dLN and ndLN at various time points. Specifically, IL10/IL19, ICOS/CD38, CD172a/MERTK and Cell cycle GEPs were up regulated in dLN at V4 compared to V2 (Figure S12A-C). From comparison with the Pan-Cancer Atlas gene sets, cytokine, chemokine, cytotoxicity, and anergy gene sets were sharply downregulated at V4 in the dLN compared to V2 (Figure S12D). At the metabolic level, the oxidative phosphorylation (OXPHOS) gene set was upregulated at V4 in both the dLN and the ndLN (Figure S12D). Projection of GEPs derived from our cNMF analysis across time points showed that GEP1 from both CD4^+^ and CD8^+^ T cells was increased in the dLN V4 compared to baseline dLN V2 (Figure S12E-F), aligned with the increased activity of *BACH2* in the dLN following immunisation. BACH2^+^ and ARHGAP15^+^ T cells displayed similar TF profiles, such as an anti-apoptotic programme, that were distinct from classical T cells, while each state maintained distinctive phenotypic and functional identities, especially Tfh.

### The quiescence T cell states showed a divergent differentiation trajectory distinct from classical T cell lineages, including Tfh

To understand trajectories of LN T cells following immunisation, the principal graph algorithm was implemented by scFates^39^, which reconstructed cell-fate lineages and inferred pseudotime. Naive T cells were defined as the root state, yielding distinct developmental trajectories for BACH2^+^, ARHGAP15^+^ and classical T cell lineages for both CD4^+^ and CD8^+^ T cells (Figure 5A and S13A). Compared to the Tfh lineage, the BACH2^+^ lineage exhibited progressive increase of *LEF1* and *TCF7* along pseudotime. Conversely, the classical lineage cells showed a steady decline of these markers (Figure 5B-C). Notably, *PTMA* was inversely correlated with the kinetics of *LEF1* and *BACH2* within the BACH2^+^ lineage, while this was steadily maintained within the conventional Tfh lineage. *ICOS* increased along the trajectory in the Tfh lineage and *GZMK* and *GZMB* dynamics in the CD8^+^ T cell lineage, confirming expression kinetics from existing literature^40^ (Figure 5B-C). Moreover, *ARHGAP15* and *UTRN* expression increased along pseudotime in ARHGAP15^+^ T cells, whereas *PTMA* expression declined in both ARHGAP15^+^ and BACH2^+^ T cell states (Figure S13B).

**Figure 5.**
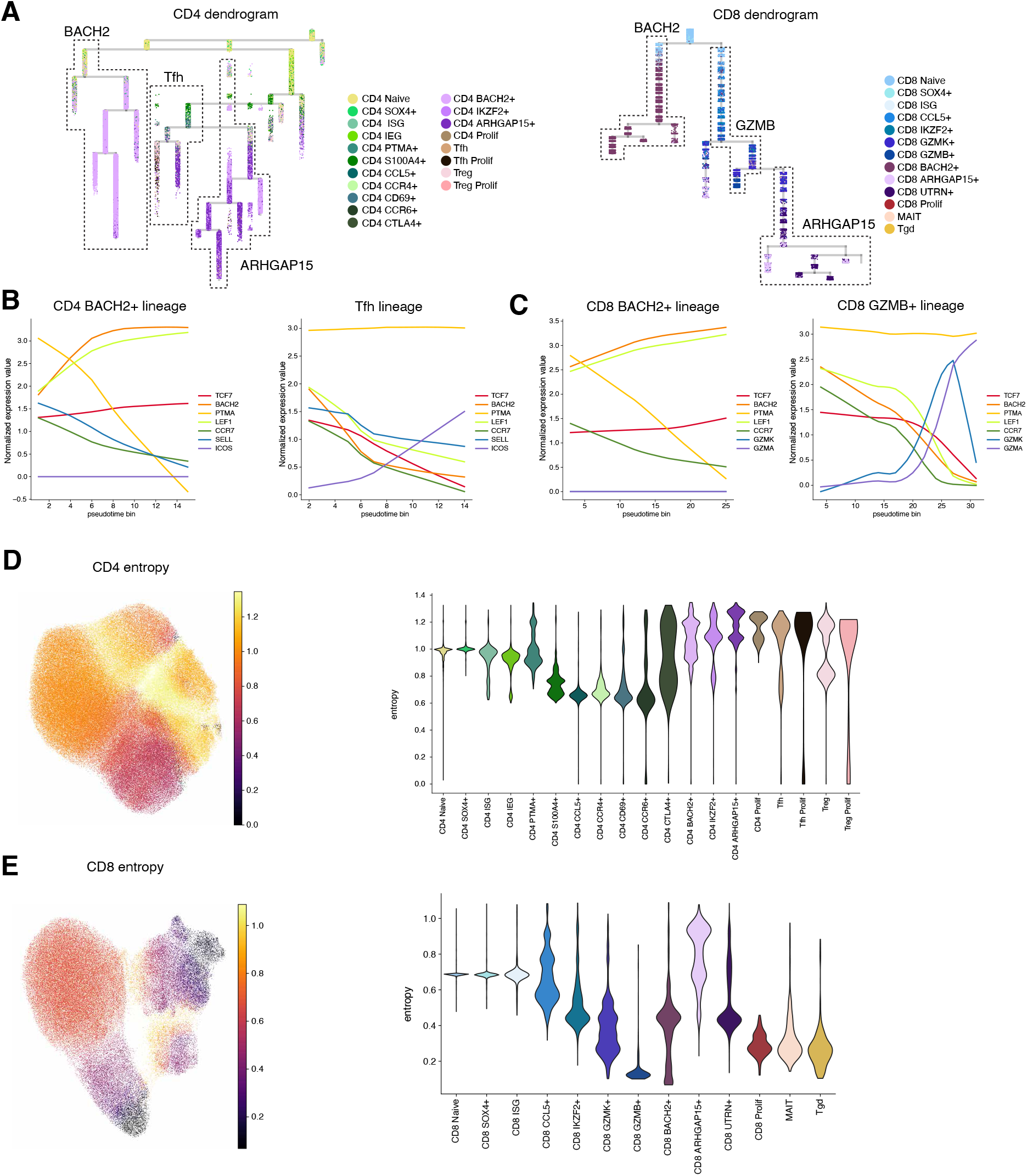
Trajectory mapping of human LN T cells. (A) Dendrogram depicting the differentiation trajectories from CD4^+^ (left) and CD8^+^ (right) T cell and colored by cell state. (B) Expression patterns of selected expressed genes along the trajectory in BACH2^+^ lineage (left) and Tfh lineage (right) from CD4^+^ T cell and (C) BACH2 lineage (left) and GZMB^+^ lineage from CD8^+^ T cell. (D) Simple embedding colored by inferred differentiation potential (left) and Violin plots of entropy distributions across cell types (right) for CD4^+^ T cell and (E) same for CD8^+^ T cell.

Using Palantir^41^ detected different Shannon entropy amongst these different LNC states (Figure 5D-E). Amongst CD4^+^ T cells, BACH2^+^, ARHGAP15^+^, proliferating and Tfh had high entropies, suggesting that these subsets reside in a less uncommitted state of lineage differentiation. Conversely, there was a steady decrease in entropy along the classical T-cell differentiation path. The entropy in dLN amongst CD8^+^ T cells peaked post immunisation in dLN at V4 (Figure S13C). Classical T cell lineages, including the Tfh and CD8+ GZMB+ lineages, followed expected gene expression along the trajectory whilst BACH2^+^ and ARHGAP15^+^ T cell states occupied distinct developmental branches that retained features of quiescence, stemness and/or entropy.

### Clonal sharing amongst different T cell subsets induced by immunisation

To elucidate the clonal relationships between populations, we next investigated the TCR repertoires induced upon immunisation. Clonal expansion analysis revealed T cells with low TCR signals within the quiescent T cell states (Figure 6A and S14A). It is possible that the lower expression of TCR by these cell states resulted from an attenuated TCR activation program that supports a quiescent, stem-like state^25^ (Figure S2C). Integrating with pseudotime results revealed that cells lacking TCR repertoires showed a sharper increase along the pseudotime trajectory, particularly within BACH2^+^ and ARHGAP15^+^ T cell lineages (Figure S14B-C). To understand the potential influence of technical artifact on these findings, TCR recovery rates per UMI counts were evaluated. Quiescent T cell states showed an upward trend of TCR recovery rates per depth, and a lower rate with equalised sequencing depth. Although lower baseline UMI sequencing depth could account for the observed cell states lacking TCR expression, a similar result was observed in the previously published study on influenza immunisation in the dLN, suggesting this finding reflects intrinsic biological differences in TCR expression across T cell states (Figure S15A-C)^28^. Further characterisation of TFs and GEPs in TCR-low and TCR-hi cells revealed an upregulation of BCL2 anti-apoptotic pathways and GEP1 in TCR-low cells, suggesting that TCR-low T cells exhibit enhanced BACH2-like characteristics (Figure S15D-E).

**Figure 6.**
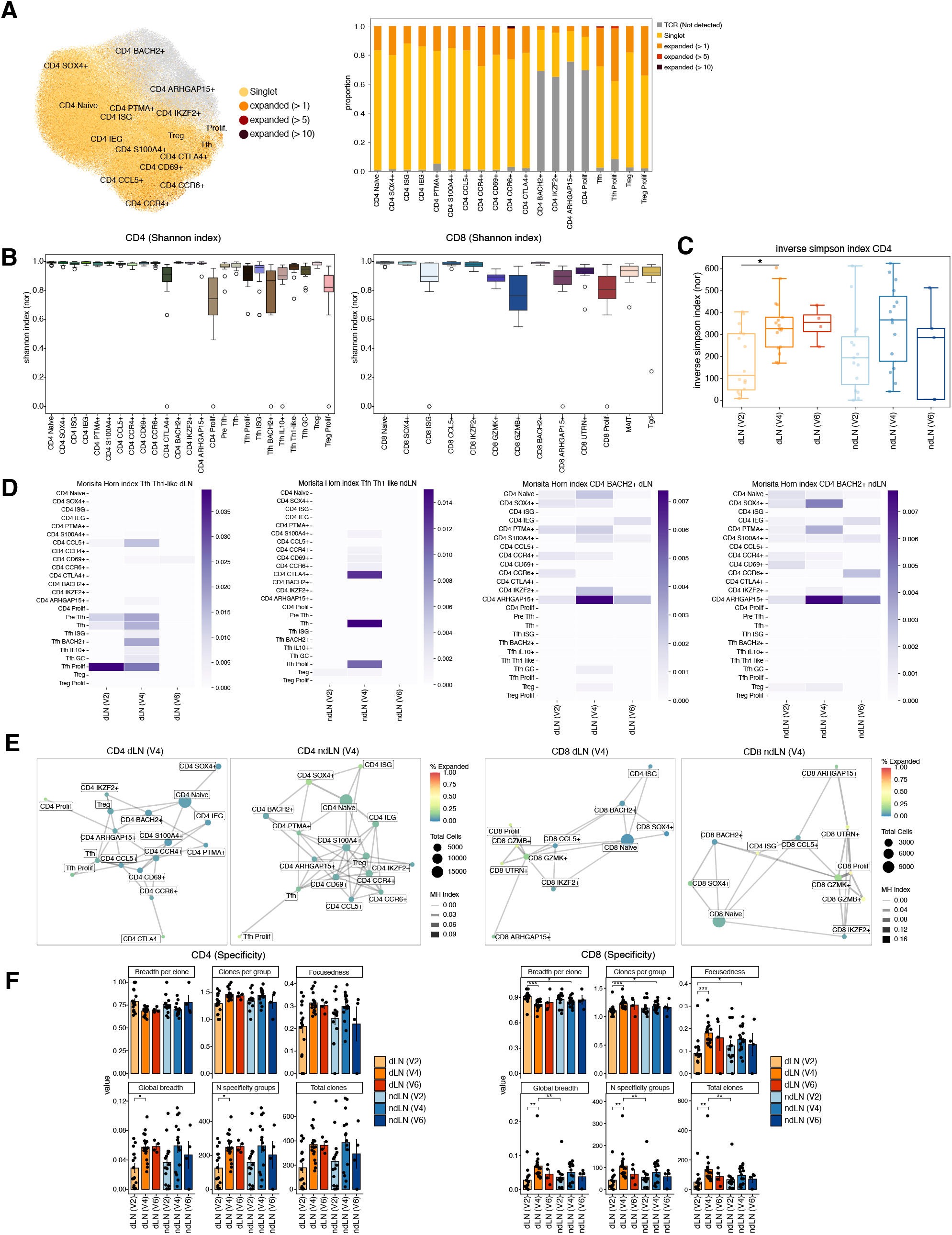
Immune receptor repertoire across cell states in human LN across sample sites and time points. (A) UMAP visualisation (left) and stacked bar plots (right) show the proportion of clonally expanded CD4^+^ T cells. (B) Normalised Shannon entropy index was estimated for each cell state in the CD4^+^ (left) and CD8^+^ (right) T cell. (C) Box plot representing normalised inverse Simpson index for a group of different time points per each donor. Statistical significance is defined using Wilcoxon rank-sum test and depicted as *p<0.05. (D) Clonal dynamics between cell-types as indicated by Morisita-Horn (MH) similarity index along the time points and sites for Th1-like Tfh in dLN (left) and ndLN (right) and same for CD4^+^ BACH2^+^ T state. (E) Shared clonal networks at V4 of CD4^+^ T cell (left) and CD8^+^ T cell (right) repertoires after filtering, where edge weights are defined by MH similarity index, node colour denotes clonality, and node size indicates number of clones. (F) Box plots denote different metrics of distinct clonotypes of specificity across sample sites and time points CD4^+^ T (left) and CD8^+^ T (right) (See more detail in Methods).

To understand TCR dynamics between cell states, we compared TCR repertoire diversity metrics. Evaluation of alpha diversity across cell types using the normalised Shannon index revealed a selective reduction in CD4^+^ proliferating, CTLA4^+^, proliferating Treg and Tfh subsets, as well as ISG, GZMK^+^, GZMB^+^, ARHGAP15^+^, proliferating within CD8^+^ lineage (Figure 6B). This lower repertoire diversity suggests the occurrence of antigen-driven clonal expansion in these subsets, further validated by a decline in Pielou’s Evenness indicating clonotypes were distributed less evenly. Cell types containing expanded clonotypes exhibited lower normalised inverse Simpson index values suggesting increased dominance of abundant clonotypes (Figure S16A-B). This clonotype dominance (by inverse Simpson index) in the draining LN increased over time in CD4^+^ T cells, whilst in CD8^+^ T cells it peaked at V4 and decreased afterwards, albeit in small sample numbers (Figure 6C and S16C).

The Morisita-Horn index was employed to compare TCR repertoire overlap dynamics between cell types across dLN and ndLN and sampling time points. Clonal overlap increased at V4 compared to V2 within the Tfh subsets, which subsequently declined at V6 in dLN (Figure S16D and S17A) and there was some overlap in Tfh subset cells in ndLN (Figure S17B). Focused analysis on subsets including Th1-like Tfh showed increased overlap with CD4^+^ CCL5^+^ and Tfh subsets in the dLN at V4 (Figure 6D). CD4^+^ BACH2^+^ T cells had increased overlap with CD4^+^ ARHGAP15^+^, suggesting shared clonal dynamics between these cell states. Amongst CD8^+^ BACH2^+^ T cells, clonal overlap was with less differentiated, early cell types in the dLN at V4 (Figure S17C). The highest degree of clonal overlap by the Morisita-Horn index between timepoints was for V2 and V4 in the dLN amongst both CD4^+^ and CD8^+^ T cells, with considerable overlap between the dLN and ndLN at V6 amongst CD8^+^ T cells (Figure S17D).

In addition to clonal relationships observed at V2, including those between Tfh and proliferating Tfh cells and between CD4^+^ CCR4^+^ and CCR6^+^ T cells, there was increased clonal connectivity between CD4^+^ naive, IEG, S100A4^+^ and BACH2^+^ T cells following immunisation (Figure 6E and S17E). There was increased clonal sharing between CD4^+^ ARHGAP15^+^, BACH2^+^ and multiple classical T cells, including CCL5^+^, Tfh, Treg, IKZF2^+^, and proliferating T-cell populations (Figure 6E). In contrast, clonal connectivities among CD8^+^ GZMK^+^ and GZMB^+^, CCL5^+^ T cells were less prominent upon immunisation, with increased clonal relationships between naive and quiescent CD8^+^ T-cell states (Figure 6E and S17E). These observations suggest increased clonal diversification of naive CD4+ T cells into multiple states with early activation and differentiation in response to immunisation.

To investigate the antigen specificity of vaccine-responsive T cells, TCR sequences were mapped to publicly available databases and analysed with ImmunoWatch^42^ (Figure S18A-B). The majority of TCRs detected in our dataset could not be assigned to known antigen-TCR pairs. Amongs TCRs with annotated specificities, global changes across cell states following immunisation were not observed. Similarly, computational predictions with public databases did not reveal strong enrichment of influenza epitopes after vaccination. CD4^+^ ISG T cells appeared induced in draining LNs following vaccination (Figure S18C). To further characterise antigen specificity, we grouped TCRs using TCRdist3^43^, which identifies clusters of TCRs predicted to recognise shared antigens. Both CD4^+^ and CD8^+^ T cells showed an increase in the number and breadth of specificity groups in draining LNs following immunisation (Figure 6F). Although inferring precise antigen specificity remains limited, these findings demonstrate coordinated TCR repertoire dynamics across distinct lymph node T cell populations following vaccination.

### Spatial co-localisation of BACH2^+^ T cell and Th1-like Tfh in the lymph node T zone

To investigate localisation and interactions of T cell states identified in response to immunisation, we first annotated B and myeloid populations (Figure S19A-B) and performed cell2location^44^ to assign cell type densities to spatial coordinates using our scRNA-seq mapped to publicly available human LN spatial data. The spatial deconvolution identified that BACH2^+^ and ARHGAP15^+^ T cells and Th1-like Tfh cells were co-localised outside of the follicular areas and within the LN T cell zone. Dendritic cell (DC) LAMP3^+^ co-located within this same T cell zone, whereas other myeloid and B cell subsets did not (Figure 7A and S19C). NMF decomposition confirmed that functionally coordinated cellular compartments clustered within their expected anatomical LN site. As expected, Tfh GC cells explicitly co-localised with B-cell GC light zone cells, whilst the rest of the Tfh subsets co-localised within the T zone; factor 6 (Figure 7B).

**Figure 7.**
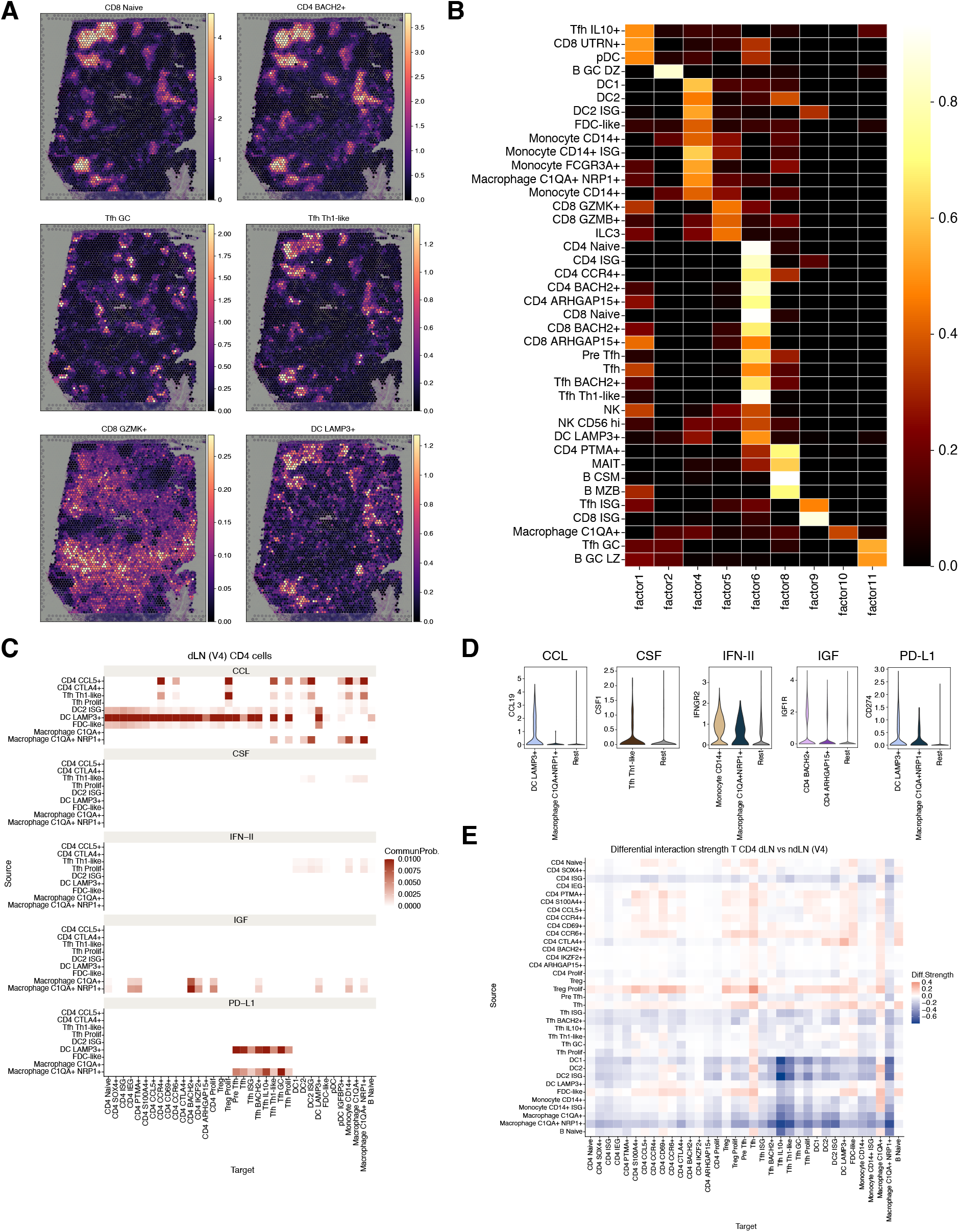
Mapping the cellular compartments and interactions of the human LN. (A) Estimated cell abundance of selected cell types by color intensity (estimated cell density) with highlighted cellular compartments. (B) Heatmap showing NMF factors, where each factor is a group of co-localised cell types. The color density represents the loading of each cell type in each factor. (C) Heatmap showing significant signaling predictions for each signaling family between each sender (ligand) and target (receptor) cell type. The color indicates the strength of signaling (interaction probability). (D) Violin plot showing expression genes of each signaling family. (E) Heatmap showing overall differential interaction strength between dLN and ndLN at V4.

Cellular communications within the LN were interrogated to understand local signaling events. There was a prominent chemokine ligand (CCL) signaling axes, especially CCL19, between DC LAMP3^+^ and T cells. Specifically, both CD4^+^ and CD8^+^ BACH2^+^ T cells interacted with macrophage C1QA^+^ NRP^+^ cells via the insulin-like growth factor (IGF) signaling axis (Figure 7C and S20A). Furthermore, a specific relationship was identified where the PD-L1 signaling axis connected DC LAMP3^+^ and macrophage C1QA^+^ NRP^+^ cells with all Tfh subsets. Specific signaling interactions between Th1-like Tfh cell and myeloid cell subsets through Type II interferon and colony stimulating factor signaling were also detected (Figure 7C-D and S20A).

We subsequently evaluated differences between dLN and ndLN at V4, and found that the CCL signaling axis operating between DC LAMP3^+^ cell type and T cells was increased within dLN, but decreased in FDC-like cells (Figure S20B). The IGF signaling axis decreased in dLN in both CD4^+^ and CD8^+^ BACH2^+^ T cells (Figure S20B-D). The strength of ligand and receptor interactions was lower in the dLN involving both CD4^+^ and CD8^+^ ISG cells, while interactions involving proliferating Treg and Tfh cells were increased at V4. Interestingly, the Tfh subset and macrophage C1QA^+^ cells had increased incoming receptor signaling within the dLN (Figure 7E and S20E). Taken together, spatial mapping onto publicly available data and cell-cell interaction analysis of the study data suggested a spatially interlaced cellular landscape in the T cell zone amongst vaccine-responsive dLN T cell subsets.

## Discussion

This study demonstrated that adjuvanted influenza vaccine induces multiple transcriptionally distinct T cell subsets in the dLN of African and Asian young adults in the first 5 days after injection, with Th-1 like Tfh and BACH2+ T cells states emerging as characteristic components of the response. We did not find major ancestrally driven differences in the T cell LN response to immunisation, which may reflect a small effect size, as this study was designed to describe LN responses rather than specifically test differences in ancestry. This finding is consistent with previous studies which found that age but not ethnicity was associated with immune responses to influenza vaccine^45,46^. Twin studies also suggested the impact of heritability on influenza vaccine response may be low, although ancestry driven changes in B cell antibody diversity genes have been detected ^47,48^.

The prominent Th-1 like Tfh responses observed, with transcriptionally unique signatures compared with Tfh GC, suggests that these cells were not located in the germinal centre at the time of sampling. Furthermore, this cell state was found in the extrafollicular regions when mapped to publicly available spatial LN data. These Tfh cells were highly activated transcriptionally by immunisation, and we and others have previously demonstrated similar induction of Tfh cells^19,49^. Clonotype dynamics also indicated a change in clonal abundance in these cells with immunisation. Whilst multiple Tfh subsets were induced by vaccination, these findings additionally pinpointed Th1-like Tfh expressing CXCR3 as strongly induced in dLN. In the blood, the circulating counterpart of this cell type has been correlated with support for B cells and antibody responses in several studies^50–52^. This population may emerge from a Th1/Tfh precursor defined by TCF1, SLAMF6, PD-1, BCL6, CXCR3 and CXCR5 as has been shown in mice^53^. Whilst a recent dLN single-cell study that mapped early influenza vaccine responses primary focused on GC Tfh and IL10^+^ Tfh kinetics, the study did not identify this distinct Th1-like Tfh cell state^28^.

The previous study used a non-adjuvanted vaccine, unlike the MF59C.1 adjuvanted aQIV used here, which raises the question as to whether the Th1-like Tfh cell state we observed is a product of adjuvant exposure in the dLN. Whilst our data indicated a phenotypical difference of CXCR3^+^ Th1-like Tfh from Tfh GC cells in the lymph node, most previous studies have focused on circulating Tfh subsets (cTfh) due to limitations in sample access. In one study, CXCR3 negative cTfh were transcriptionally similar to tonsil GC Tfh and associated with HIV broadly neutralizing antibody development, and CXCR3+ cTfh had a Th1-like phenotype with secretion of interferon-gamma in HIV infection^54^. In SARS-CoV-2 infection, however circulating Th-1 like CXCR3+ Tfh cells were associated with higher antibody titers post immunisation and CXCR3-cells were not^55^. Differences in GC Tfh and Th1-like Tfh may be due to *TOX2* expression; we observed less *TOX2* expression in Th1-like Tfh cells. High *TOX2* expression was almost exclusive to TCR-stimulated Tfh GC cells, which maintained their transcriptional programme inhibiting spontaneous conversion into Th1-like cells^31^. Our data raise the possibility that Th1-like Tfh in the lymph node are involved in the early dLN response to the recall antigens in the adjuvanted seasonal influenza vaccine, and that this is, initially, both spatially and temporally distinct from the GC response.

Amongst the other T cell states induced following immunisation, some subsets exhibited a transcriptional programme marked by the expression of *BACH2*, *ARHGAP15*, *IKZF2* and *UTRN.* These were transcriptionally distinct from GC responses, and different from classical T cell lineages, such as naive, central and effector memory T cells. These states had low ribosomal counts, which is a feature for maintaining quiescence^24,56^. T cells expressing *BACH2* were surprisingly dominant within these vaccine-induced cell states in the dLN. BACH2 acts as a regulator of immune cell differentiation, a TF well described in B cells, but less so for T cells^57^. While its role in human LN function has not been well described, what is known is that its expression induces quiescence and exhausted progenitors. This occurs with induction of stem-like genes including *LEF1* and *TCF7* that restrain terminal differentiation, and increase of the pro-survival factor *BCL2*, with attenuation of AP-1 factor genes and TCR signaling^16,58–60^. Recently, modulation of BACH2 expression was shown to balance the stemness or exhaustion phenotypes of CAR T cells^58^. Our hypothesis that BACH2 expression is associated with long term T cell memory formation is supported by studies of HIV infection. HIV exploits BACH2-induced T cell states to form long-lived memory T cells, promoting seeding and maintenance of HIV infection as the viral reservoir^61^. Epigenetically, BACH2 enhances chromatin accessibility at loci linked to stemness-associated TFs and anti-apoptosis gene expression^25^. This suggests BACH2 as a potential candidate for a molecular rheostat governing T cell fate determination in the early immune response to immunisation in the human dLN.

Another notable feature of these vaccine-induced T cell states was high expression of chemokine receptor proteins, including CX3CR1 and CXCR3, suggesting migratory potential. Specifically, CX3CR1 and CXCR3 serve as key receptors for fractalkine (CX3CL1) and CXCL9-11, respectively; important chemokines that drive cellular positioning^62^. Expression of CXCR3 was rapidly upregulated and remained high on antigen-specific T cells in mouse dLN with a role in LN positioning to promote antigen recognition^63^. In blood-borne memory T cells, viral infection induced a stable population of CX3CR1int memory CD8+ T cells which unique phenotypic, homeostatic, and migratory properties^64^. Expression of CXCR3 on vaccine induced T cells may thus be an important component of cellular positioning within the dLN, which was notable with our use of an adjuvanted seasonal influenza vaccine.

Whilst we were unable to directly characterise the spatial positioning of these interesting T cell subsets in study participants, comparison with publicly available data corroborated the hypothesis that these cells were largely outside the GC and possibly extrafollicular at the time of sampling. BACH2^+^ T cells and Th1-like Tfh were co-located in the T cell zone, while Tfh GC colocalised with B GC LZ cells through mapping to publicly available spatial LN data. Migratory DC LAMP3^+^ cells were also colocalised within the T cell zone and showed interactions with these cell types in our data. Migratory DCs were required for clonal selection and expansion of CD4^+^ T cells after subcutaneous immunisation and this directional migration of both DCs and T cells into the T cell zone was mediated by the chemokine receptor *CCR7*^65,66^. This ligand–receptor communication was also confirmed spatially using CellNEST with T cell zones as the primary location for CCL19–CCR7 axis^67^. Ligand-receptor communication also revealed enrichment of IGF-1 signaling axis between BACH2^+^ T cells and macrophage subsets, which may be functionally relevant given macrophage-derived IGF-1 deficiency is associated with lower influenza vaccine specific antibody in mice^68^.

### Limitations

Due to the small sample size, we were unable to directly test for vaccine related differences in the immune response arising from different ancestral genotypes. The ancestral variations identified in this study provide a foundation of hypothesis-generating insights, highlighting the need to further investigate the functional immune effects of ancestrally driven genetic diversity in T cells. Given the challenge of running such studies at scale, this could be achieved computationally using the data presented here, which forms a curated on-line resource, in meta-analyses with other similar studies. Our differential abundance analyses were broadly consistent with observations in cohort 1 that used MASC to test differences in abundance, albeit with some differences in cell subset annotation^19^. The studies tested slightly different time points as allowed by the protocol, with a greater spread of timepoints in cohort 1. Both studies identified Tfh as dominant in the dLN post immunisation. The current study additionally identified a transcriptional quiescent cell state but was unable to determine the mechanistic consequence of BACH2 induction in CD4^+^ and CD8^+^ T cells. Whilst we could not entirely control for the possibility of technical artifacts due to low TCR recovery rate, based on the available literature on BACH2 function, we hypothesise a role in the induction of T cell immune memory that could be tested in future work. Ultimately, larger follow up studies with repeated follow-up timepoints will add precision to our findings on the kinetics of the T cell immune response in the dLN. Whilst a goal of the project was to identify influenza vaccine-specific TCR, these epitopes, from an ancestrally diverse cohort, remain poorly represented in current reference databases. This highlights an opportunity for further research to define the epitopes recognised by these clonotypes, to which these data will also contribute.

This study demonstrated an early T cell immune response in the dLN of young adults who self-identify as having African or Asian ancestry in response to adjuvated influenza vaccine, with several defining features. These included both activated support of the antibody response by Th1-like Tfh and expression of the TF BACH2. Given these cells were transcriptionally unique, and highly dominant in the dLN, both findings may be important for subsequent immune memory formation and therefore functionally relevant features of the vaccine-induced immune response. Whilst the function of Tfh in the dLN has been well described in animal models, the function of this BACH2^+^ T cell state is unknown. This vaccine-induced T cell population actively engaged a highly specialised transcriptional signature optimised for cellular survival and long-term persistence and may represent an important component of the early dLN extrafollicular T cell response to immunisation. We present our data as a curated on-line resource to contribute to addressing paucity of knowledge in lymph node T cell function, and the under representation of these ancestries in immunity research.

## Methods

### Clinical study and sample collection

The clinical study, participant eligibility and enrolment have been described previously (Siu et al.) In brief, LEGACY01 (ISRCTN13657999) study was a single-site interventional non-randomised open label experimental medicine clinical research study over two influenza seasons (2022–2023 and 2023–2024). The study was run at the National Institute for Health and Care Research (NIHR) Imperial Clinical Research Facility, West London, UK and approved by the London - Central Research Ethics Committee (ref: 22/LO/0343). We report here data from the second season cohort. Participants were healthy adults aged ≥18 years and ≤55 years (median (IQR) 26–40 years old) of several different self-declared ethnicities of African and Asian ancestry. In cohort 2, n=17 participants received aQIV by intramuscular injection into the deltoid muscle of the arm. Arm selection for injection (left or right) was by participant choice and was recorded by the study team. We collected draining and non-draining axillary LN cells from 17 participants at 3 time points by US guided FNA of left and right axillae: pre-vaccination (V2) and post-vaccination at day 5 (V4) and, optionally at 6 weeks (V6) with adjuvanted influenza vaccine (aQIV) under US-guided FNA of dLN and ndLN. In two donors (LEG1048 and LEG1050), the post-vaccination FNA (V4) was at 13 and 12 days respectively. Up to five participants were given the option to attend an additional US-guided FNA at 6-10 weeks post immunisation at visit 6 (V6), based on staff availability to perform the procedure. Samples processing was performed as described previously^14^. There was no safety objective of the study. To ensure the ethical conduct of the study, safety oversight was conducted by the clinical team with non-serious adverse events reviewed up to 5 days post FNA and 28 days post immunisation, and serious adverse events throughout the study, according to the protocol (LEGACY01 Protocol v3.1 06Jul23) and in line with Sponsor requirements for an experimental medicine study (Supplementary Material).

### Study vaccination

Participants were immunised with the adjuvanted quadrivalent influenza vaccine (aQIV) (CSL Seqirus), by intramuscular injection into the deltoid region of the non-dominant arm. For cohort 2, the 2023-2024 winter season vaccine was used, which contained, 15 micrograms each of the following haemagglutinins, A/Victoria/4897/2022(H1N1)pdm09-like strain, A/Darwin/9/2021 (H3N2)-like strain, B/Austria/1359417/2021-like strain and B/Phuket/3073/2013-like strain. The adjuvant was MF59C.1 as previously described^19^.

### HLA genotyping and ancestry proportion estimates

DNA was isolated from whole blood using the QIAamp DNA Blood Midi Kit according to manufacturer’s instructions. Genotyping was performed using the Infinium Global Screening Array-24 v3.0 (Illumina) by the Genome Centre, Queen Mary University of London according to manufacturer’s recommendations. HLA typing up to four digits, was performed by the sequencing/typing facility at the MRC Weatherall Institute of Molecular Medicine, University of Oxford. The unprocessed genotyping data consisted of 17 volunteers and 654,027 variants prior to filtering and QC using PLINK^69,70^. Filtering and QC steps included: (1) retaining only autosomal variants, (2) removing loci where 99.9% of genotypes are missing, (3) removing SNPs with minor allele frequency of >1%, and (4) excluding variants with one or more multi-character allele code. The final post-QC array data consisted of 336,767 variants. The reference dataset^71^ combines genotypes (n=3433) with known ancestry from the 1000 Genomes Project^20^ and Human Genome Diversity Project (HGDP)^21^. Intersecting variants between our data and reference data for downstream analysis was determined by the following: (1) exclude non-unique SNPs, (2) exclude A-T or G-C SNPs, and (3) aligned position mismatches and allele flips. The combined data was pruned for linkage disequilibrium using a window size of 50 variants, window shift at each step of 5 variants and r^2 threshold of 0.2 (PLINK setting: “--indep-pairwise 50 5 0.2”). The final pruned combined dataset included 84,543 variants. The program ADMIXTURE^22^ was used to estimate per-individual ancestry populations in a panel of 3433 reference individuals representing African, European, East Asian, and American ancestries^71^. The optimum number of ancestry populations (K) was chosen based on five-fold cross-validation for each K in the set of 1–9. K = 7 was chosen based on no additional improvement in terms of the cross-validation error for larger K (Figure S1A). The population allele frequencies estimated from the analysis of reference samples were fixed as parameters so that the LEGACY samples could be projected into the admixture model to obtain ancestry proportion estimates.

### Library preparation and sequencing of scRNA-seq and CITE-seq samples

Longitudinal samples from 17 donors at three time-points were processed using the CITE-seq method. Samples were stained with unique hashtag oligo-conjugated antibodies (HTO) for sample pooling and other antibody-derived tag antibodies (ADTs) targeting specific surface-expressed proteins using a manually constructed pool of 63 TotalSeq-C antibodies or the 137 target TotalSeq-C Human Universal Cocktail V1.0 (all BioLegend), before cell partitioning on the Chromium Controller. Around 60,000 cells were loaded into each well of GEM-X Chromium chips G, and GEX and protein (HTO/ADT) libraries were constructed as previously described^14^. After loading onto the channels of a Chromium chip (10× Genomics), cDNA synthesis, amplification and sequencing libraries were generated using Single Cell 5ʹ Reagent (v3) kit. Also, TCRαβ and BCR paired VDJ libraries were prepared from samples made with the 5ʹ Reagent Kit. All libraries were sequenced on NovaSeq X instrument.

### Alignment and quantification of scRNA-seq data

Reads from scRNA-seq and VDJ libraries were alignment to the 10x Genomics human reference genome GRCh38 (2020-A and vdj-GRCh38-alts-ensembl-7.1.0), followed by cell calling, transcript quantification and sequence assembly and paired clonotype calling on the V(D)J libraries using the Cell Ranger multi (v.8.0.1; 10x Genomics) with default parameters using Nextflow (v.24.04.2). Cell Ranger filtered count matrices were used for downstream analysis.

### Demultiplexing

Hashtag data were demultiplexed using HashSolo^72^, with default parameters. However, some data included a mixture of hashtag and non-hashtag samples (Table S4). To resolve the mixing issue, we performed genotype-based demultiplexing with Souporcell (v.2.5)^73^. Briefly, variant calling was then performed using the BAM file and filtered cell barcodes using the hg38 assembly as the reference genome without prior donor genotype information. The value of *k*, representing the expected number of genotypes in each mixed sample, was assigned manually according to the specific mixing condition. After genotyping, clusters were linked back to their donor-of-origin genotypes across all mixed samples using the Souporcell shared_samples.py script. Based on the lowest-distance cluster matches, each cluster in one sample was matched to the corresponding cluster in another sample and assigned to a mixed donor. Cells classified as negative or doublets by HashSolo, as well as genotype doublets and unassigned cells, were excluded from downstream analyses.

### Quality control, filtering, and preprocessing of the data

Data were analysed using Scanpy (v.1.11.0)^74^ following their standard recommended pipelines. Low-quality cells (1) expressing fewer than 500 genes, (2) mitochondrial content higher than 20%, (3) expressing more than 100k UMI counts were removed from downstream analysis. For doublet removal, we applied Scrublet (v.0.2.3)^75^ to each sample as described previously^76^. Briefly, we performed a two-step diffusion doublet approach to propagate Scrublet scores to similar barcodes, followed by Bonferroni correction for multiple testing. Barcodes estimated as potential doublets and doublet-dominated clusters were excluded. Preprocessing steps, including normalisation and scaling, were performed independently for each analysis using the normalise_total and scale functions in Scanpy. Following PCA, we computed the neighborhood graph, generated a uniform manifold approximation and projection (UMAP) visualisation and performed Leiden clustering per each sample. For broad annotation, we applied a CellTypist (v.1.6.3)^77^ approach with pre-trained model (Healthy_COVID19_PBMC) and a customised LEGACY01 Cohort1 model with default settings. For each sample, cells were retained when both models assigned consistent broad immune categories. Cells with discordant lineage assignments between the two models were flagged for further investigation to determine whether they should be included or excluded from downstream analysis.

### Integration and batch correction of datasets

After removal of lowQC cells and doublets, we integrated all samples by an MrVI^78^ model from scvi-tools (v.1.3.1) on gene expression data using 5,000 highly variable genes selected with the seurat_v3 option in Scanpy. All the parameters were kept as default, with sample as sample key and previous annotation as label key and trained 500 epoch. Louvain clustering and UMAP embeddings were generated using MrVI latent output. After batch correction and T cell lineage separation, we repeated MrVI integration on T cells only, as described above and finally repeated on CD4, CD8 T cells and NK/ILC, respectively. Myeloid, B-lineage and Tfh subset were batch-corrected using Harmony^79^ (v.0.0.6) with sample as a covariate, utilising principal components. Similarly, louvain clustering and UMAP embeddings were generated using Harmony latent output.

### Sub-clustering and cell type annotation

Cell-type annotations were annotated using well-characterised marker genes by identifying cluster-specific marker genes using rank_genes_groups with the Wilcoxon test and CITE-seq surface protein expression. To align annotations with previous findings, we utilised customised LEGACY01 Cohort 1^19^ CellTypist labels and further refined with cluster-specific marker gene expression. Within each Cluster, concordant between marker gene and CellTypist assignments were retained, whereas discordant marker gene and CellTypist annotations were manually re-annotated based on marker genes. CITE-seq antibody-derived tag counts were separated from the gene expression matrix and normalised using centered log-ratio (CLR) transformation. The resulting values represent the log-transformed abundance of each protein relative to other proteins measured within the same cell. Dot plots and violin plots were produced in Scanpy internal functions. Unless otherwise stated, displayed gene expression values were log-normalised in the preprocessing section. For dot plots spanning multiple datasets, each dataset was independently log-normalised, variance-scaled, and min-max standardised to a 0–1 range.

### Estimation of variation contributions of each feature

Variance component models were fitted using the variancePartition^80^(v.1.32.5) to estimate the sources of variation from a list of covariates for each feature in the pseudobulk transcriptomic data. We first aggregated data across individual and broad cell types using decoupleR^81^ (v.2.1.1). Then, genes with count more than 10 and at least 3 of samples were kept and normalised using vst function from DESeq2^82^ (v.1.42.1). Ancestry was assigned based on the largest ancestry proportion into 4 groups. The normalised expression was used to model with days as a fixed effect while sex, annotation and ancestry were modeled as a random effect

### Comparison with public human LN data

To assess a quantitative measure of the similarity with the public QIV LN dataset^28^, first we integrated our data with the reference dataset and performed batch correction using Harmony as described. Similarity between annotations or clusters in the public QIV LN dataset and our annotations was then quantified using cosine distance, calculated with the Distance function implemented in the Pertpy^83^ (v.0.10.0).

### Differential abundance testing

Bayesian hierarchical beta-binomial models were used to assess the differential abundance of cell states. The number of cells assigned to that state was modelled as the response, with the total number of cells per sample included as the beta-binomial denominator. Models were fitted separately for LN status comparisons (dLN versus ndLN) and timepoint, with age and BMI included as fixed-effect covariates. Individual participant and batch effect were included as random effects to account for repeated sampling and technical variation. Weakly informative priors were specified for both fixed and random effects. Posterior samples were obtained using Hamiltonian Monte Carlo with 9,000 iterations, including 4,500 warm-up iterations, across eight chains. Posterior odds ratio estimates were summarised as posterior medians with 95% credible intervals. Model estimates were taken as significant where the 95% credible interval excluded 1, and no p-values are calculated. Model convergence was assessed using R-hat metric, and model fit was evaluated by visual posterior predictive checks. Models were implemented in R using the brms^84^ (v.2.23) with cmdstanr (v.0.9.0) and cmdstan (v2.39.0) serving as interfaces to the Stan probabilistic programming language^85^.

### Differential gene expression analysis and gene expression program pathway

To prevent individual samples with a high number of cells dominating downstream analysis, cells were downsampled to around the median cell number per sample across each cell type. To identify differentially regulated genes among conditions, we performed empirical Bayes quasi-likelihood F-tests (QLF) including the cellular detection rate (the fraction of detected genes per cell) and sex as covariates using edgeR^86^ (v.3.28.1). A log2-fold change (FC) greater than 0.5 and a false discovery rate (FDR) less than 0.05 were considered significant. P-values were corrected for multiple testing using the Benjamini-Hochberg approach. Number of DEGs and cell counts were modeled using a negative binomial generalised linear mixed model using the brms^84^ (v.2.23). The model evaluated fixed effects for lymph node draining status and DEG direction and random intercepts for cell type. Cell counts were log-transformed in the model. Subsequently, the identified differentially expressed genes per condition underwent GSEA. First, genes were ranked according to decreasing log fold change and performed GSEA using prerank function from GSEApy^87^ (v.1.10.0) against the GO 2025 database and Reactome Pathways 2024 database^29^. Pathway programs were retained programs with FDR-adjusted p-value < 0.05 and NES (Normalised Enrichment Score) > 1

### T cell gene expression programs

T cell GEPs were obtained by querying the anndata against StarCAT^36^ (v.1.0.9, TCAT.V1), which contains GEPs derived from cNMF of 1.7 million T cells from 700 individuals across 38 tissues. We subsequently applied a non-negative least squares (NNLS) (via the starCAT pipeline) to map these reference programs onto our data. Using GEP usage profiles as features, we averaged GEP usage across patients in each group, and compared between conditions e.g. dLN and ndLN in different times or each cell state. From Pan-cancer T cell atlas^37^, the paper curated gene signatures: defined 14 transcriptional states from CD8^+^ and 12 different CD4^+^ T cell states. From the list of genes from each state, we utilised score_genes function and calculated expression. Then, as above, we averaged GEP usage across patients in each group and compared between conditions e.g. dLN and ndLN in different times or each cell type.

### TF regulatory analysis

Transcription factor (TF) activities were estimated using the decoupleR^81^ framework (v.2.1.1). Briefly, we loaded CollecTRI, a curated collection of TFs and their target genes. Then, TF activities were estimated by applying a univariate linear model (ulm) to assess the coordinated differential enrichment of pre-defined downstream target genes.

### Identify in house gene expression programs using cNMF

To identify de novo transcriptional programs across the dataset, cNMF algorithm (v.1.7.0) was performed using raw expression counts as input^36^. Briefly, cNMF algorithm decomposes the single-cell expression matrix into two distinct matrices: a gene-weight matrix defining the transcriptional composition of each program, and a cell-activity matrix representing the relative contribution of each program within individual cells. The analysis was performed independently for the broad CD4 and CD8 T cell lineages using 2,000 highly variable genes and Tfh subset using 1,000 highly variable genes across 20 iterations. The optimal number of programs k was selected by the point where the highest stability (silhouette score) and the lowest error (Frobenius reconstruction error). Based on these criteria, k = 9, 6, and 9 were chosen for CD4, CD8 T cells and Tfh, respectively. To assign functional identities, each program was annotated using over-representation analysis of its top 100 ranked genes against GO 2025 database and Reactome^29^ pathway 2024 using enrichr function from GSEApy^87^ (v.1.1.10).

### Mapping cell annotations to public LN visium data

To map cell types identified by scRNA-seq in the profiled public spatial transcriptomics slides, the cell2location^44^ (v.0.1.5) method was used and followed the tutorial of public LN. After filtered lowly expressed genes (cell_count_ cutoff = 5, cell_percentage_cutoff2 = 0.03, nonz_mean_cutoff = 1.12), reference signatures were estimated using a negative binomial regression model with batch as sample. Spatial mapping was implemented with hyperparameters (N_cells_per_location = 30 and detection_alpha = 20) and training was done with max_epochs = 30000. To identify spatial colocalizing of cell types, NMF was used on the matrix of estimated cell-type abundances. The number of factors was set at 11.

### Differentiation Trajectory Analysis

Trajectory inference and pseudo-temporal ordering were performed using scFates^39^ (v.1.1.1) and Palantir^41^ (v.1.4.4) on the CD4 and CD8 T cell lineages. To resolve branching trajectories, a principal tree structure was fitted to the multiscale diffusion space using the SimplePPT algorithm implemented in scFates. Cells were softly assigned to neighboring nodes along the principal curve. Within each lineage, a Naive T-cell cluster was manually designated as the root state. Pseudotime scoring and a developmental dendrogram were subsequently calculated along the resolved graph branches using pseudotime and dendrogram function. Expression gene pattern along the pseudotime was done by binned and fitted by generalised LOESS regression from seaborn plot. Palantir standard workflow was used for analysis of entropy using default parameters using MrVI latent. The start and terminal cells were specified manually. The entropy is quantified by Shannon entropy for each cell which explains differentiation potential.

### Ligand-Receptor Pair interaction analysis

The significant cell-cell interaction was evaluated using CellChat^88^ (v.2.0.0.9001) with human reference database. The normalised expression matrix was used for createCellChat function with default parameters and then preprocessed via the OverExpressedGene function. For each condition, the cell communication networks were aggregated via the aggregateNet function. Probability values were extracted from the CellChat object and figures were created by ggplot heatmaps. The cell types with zero probability values were assigned to rest groups.

### Single-cell TCR sequencing analysis

For single-cell TCR analysis, the standard pipeline of Scirpy^89^ (v.0.22.3) was implemented according to the standard tutorial. Briefly, we identified the immune cell-receptor composition of each cell and filtered out those without a single pair of productive TCRs defined using scirpy.pp.index_chains and scirpy.tl.chain_qc functions. We defined the clonotype using define_clonotypes function with parameters; receptor_arms = all, dual_ir = primary_only. Alpha diversity metrics were analysed with alpha_diversity function with metrics; shannon, inv_simpson, pielou_e, and chao1 and divided by log normalised cell number per condition except pielou_e. To examine clonal overlaps of TCR repertoires, Morisita-Horn index was calculated based on the script from pyTCR^90^ overlap_analysis (github.com/Mangul-Lab-USC/pyTCR/) and visualised as heatmaps. For shared clonal network analysis and visualisation, the R package igraph was used to construct weighted, undirected T-cell networks, with edge weights defined by Morisita’s clone overlap index, node colour denoting clonality, and node size denoting number of clones. We also filtered out edges with very low clonal overlap based on the distribution of Morisita’s index. Networks were visualised using the R package ggraph with graphs laid out for visualisation using a force-directed Fruchterman-Reingold layout.

### Breadth of antigen-specific clusters

Similarity networks were generated by connecting clonotypes with a TCRdist3^43^ (v.0.2.2) distance ≤18, a commonly used threshold representing highly similar TCRs predicted to recognise related antigens. To avoid excluding clonotypes with only a few close neighbours, clones connected by distances ≤40 were first retained before constructing the final network using the more stringent distance threshold (≤18). Edges were weighted according to inverse TCR distance, and undirected graphs were constructed using igraph. Network layouts were generated using the Kamada–Kawai algorithm, with node size proportional to clonal expansion and node composition representing the distribution of each clonotype across sampling conditions.

Connected components within the similarity network were defined as putative antigen specificity groups, representing clusters of TCRs predicted to recognise related epitopes. Antigen annotations from public reference databases were projected onto the network to visualise known antigen-specific clonotypes and evaluate the inferred specificity groups.

To quantify repertoire diversification, we calculated several complementary network metrics. Global breadth was defined as the proportion of all inferred specificity groups represented within a given sample, providing a measure of the overall diversity of predicted antigen recognition. Breadth per clonotype represents the number of specificity groups relative to the total number of clonotypes, whereas focusedness (1 − breadth per clonotype) reflects the degree to which the repertoire is concentrated on fewer antigen specificity groups. Clonotypes per specificity group measures the average number of clonotypes contributing to each predicted antigen specificity, providing an estimate of clonal expansion within antigen-specific responses. Differences between vaccination time points and anatomical compartments were assessed using pairwise Wilcoxon rank-sum tests with Benjamini– Hochberg correction for multiple testing.

### Mapping to public database

Publicly available antigen–TCR reference databases, including the VDJdb and McPAS-TCR accessed through immunarch^91^ (v.0.10.3), were integrated to annotate TCR clonotypes with known antigen specificities. Human TRB clonotypes were matched to reference databases using identical CDR3β amino acid sequences. Antigen and disease annotations from both databases were harmonised into unified pathogen categories (including Influenza, CMV, EBV, SARS-CoV-2, HIV, and others). Antigen-associated clonotypes were subsequently quantified across transcriptionally defined T-cell states, vaccination time points, and anatomical compartments to assess longitudinal changes in antigen-specific clonal composition following vaccination. TCR similarity networks were constructed separately for CD4⁺ and CD8⁺ T cells using TCRdist3-derived pairwise distance matrices calculated from TCRβ sequences. Clone-level distance matrices were imported into R, and harmonised with metadata.

### TCR recovery rate and transcriptional depth analysis

To identify TCR recovery rate across T cell states, we evaluated TCR capture per transcription depth. First, each cell state was binned based on total UMI counts and measured TCR recovery rate by number of cells with TCR divided by total number of cells and plotted in lineplot. We also subsampled each state to equalised transcription depth to 1000 and compared TCR recovery rate.

### Flow cytometry

Cryopreserved single-cell suspensions remaining after preparation for the single-cell sequencing experiments were used for spectral flow cytometry. Cells were incubated with Fc receptor blocking solution (Table S5) for 10 min at room temperature. Cells were then stained with 50 μL of surface antibody cocktail containing a viability dye (Table S6) for 1 h at 4°C. Following staining, cells were washed twice with FACS buffer (PBS supplemented with 2.5% (v/v) fetal calf serum (FCS) (Merck, Cat No F7524) and 2 mM EDTA (Merck, Cat No. EDS-100g)).

Cells were fixed and permeabilised using the BD Cytofix/Cytoperm™ Fixation/Permeabilisation Kit (BD Biosciences, CA, USA, Cat No. 554714) according to the manufacturer’s instructions. After permeabilisation, cells were incubated with Fc receptor blocking solution for 10 min at room temperature, followed by staining with the intracellular antibody cocktail (Table S7) overnight at 4°C. The following day, cells were washed with 1× Perm/Wash buffer and resuspended in FACS buffer for acquisition. Data were acquired on a 5-laser Cytek Aurora spectral flow cytometer (Cytek Biosciences, Fremont, CA, USA) and analysed by FlowJo (v10.10) (BD Life Sciences). Details of the antibodies, reagents, instruments, and software used are provided in STAR methods.

## Data and code availability

At the time of peer-reviewed publication, all analysis codes will be made available on GitHub; De-identified individual participant data that underlie the results presented in this article are available through the accompanying supplementary data. User-friendly access to single cell data from this study will be enabled by CELLxGENE. Requests for data sharing should be directed to the corresponding authors.

## Acknowledgments

We thank the participants and the PPIE members of the LEGACY committee for their support of this work. This research was conducted at the NIHR Imperial Clinical Research Facility, London UK, and supported by NIHR Imperial and Oxford Biomedical Research Centre and the Single-Cell and Spatial Genomics Facility at the Kennedy Institute of Rheumatology, University of Oxford. JHY S is supported by Wellcome Trust Early Career Award (226938/Z/23/Z). KMP is supported by an UKRI/MRC Clinician Scientist Research Fellowship (MR/W024977/1). This study was made possible in part by grant number 2021-239944 from the Chan Zuckerberg Initiative DAF, an advised fund of Silicon Valley Community Foundation.

## Declaration of generative AI and AI-assisted technologies

During the writing the manuscript, the author used Generative AI (Gemini and ChatGPT) to check grammar and phasing of statements to deliver clear messages and improve readability. The authors revised as needed and take full responsibility for the publication.

## References

1. Petrovski, S., and Goldstein, D.B. (2016). Unequal representation of genetic variation across ancestry groups creates healthcare inequality in the application of precision medicine. Genome Biol 17, 157. 10.1186/s13059-016-1016-y.

2. Kwiatkowska, K.M., Mkindi, C.G., and Nielsen, C.M. (2022). Human lymphoid tissue sampling for vaccinology. Front. Immunol. 13, 1045529. 10.3389/fimmu.2022.1045529.

3. Moysi, E., Paris, R.M., Le Grand, R., Koup, R.A., and Petrovas, C. (2022). Human lymph node immune dynamics as driver of vaccine efficacy: an understudied aspect of immune responses. Expert Review of Vaccines 21, 633–644. 10.1080/14760584.2022.2045198.

4. Kosaji, N., Zehra, B., Nassir, N., Tambi, R., Orszulak, A.R., Lim, E.T., Berdiev, B.K., Woodbury-Smith, M., and Uddin, M. (2023). Lack of ethnic diversity in single-cell transcriptomics hinders cell type detection and precision medicine inclusivity. Med 4, 217–219. 10.1016/j.medj.2023.03.002.

5. Amit, I., Ardlie, K., Arzuaga, F., Awandare, G., Bader, G., Bernier, A., Carninci, P., Donnelly, S., Eils, R., Forrest, A.R.R., et al. (2024). The commitment of the human cell atlas to humanity. Nat Commun 15, 10019. 10.1038/s41467-024-54306-x.

6. Hoang Nguyen, K.H., Le, N.V., Nguyen, P.H., Nguyen, H.H.T., Hoang, D.M., and Huynh, C.D. (2025). Human immune system: Exploring diversity across individuals and populations. Heliyon 11, e41836. 10.1016/j.heliyon.2025.e41836.

7. Peng, K., Safonova, Y., Shugay, M., Popejoy, A.B., Rodriguez, O.L., Breden, F., Brodin, P., Burkhardt, A.M., Bustamante, C., Cao-Lormeau, V.-M., et al. (2021). Diversity in immunogenomics: the value and the challenge. Nat Methods 18, 588–591. 10.1038/s41592-021-01169-5.

8. Baumgarth, N. (2025). Extrafollicular B Cell Responses—Is One Tent Big Enough? Immunological Reviews 336, e70066. 10.1111/imr.70066.

9. Elsner, R.A., and Shlomchik, M.J. (2020). Germinal Center and Extrafollicular B Cell Responses in Vaccination, Immunity, and Autoimmunity. Immunity 53, 1136–1150. 10.1016/j.immuni.2020.11.006.

10. Dhenni, R., and Phan, T.G. (2020). The geography of memory B cell reactivation in vaccine-induced immunity and in autoimmune disease relapses. Immunological Reviews 296, 62–86. 10.1111/imr.12862.

11. Huang, Q., Wu, X., Wang, Z., Chen, X., Wang, L., Lu, Y., Xiong, D., Liu, Q., Tian, Y., Lin, H., et al. (2022). The primordial differentiation of tumor-specific memory CD8+ T cells as bona fide responders to PD-1/PD-L1 blockade in draining lymph nodes. Cell 185, 4049–4066.e25. 10.1016/j.cell.2022.09.020.

12. Black, C.L., O’Halloran, A., Hung, M.-C., Srivastav, A., Lu, P., Garg, S., Jhung, M., Fry, A., Jatlaoui, T.C., Davenport, E., et al. (2022). *Vital Signs:* Influenza Hospitalizations and Vaccination Coverage by Race and Ethnicity—United States, 2009–10 Through 2021–22 Influenza Seasons. MMWR Morb. Mortal. Wkly. Rep. 71, 1366–1373. 10.15585/mmwr.mm7143e1.

13. Irving, S.A., Groom, H.C., Belongia, E.A., Crane, B., Daley, M.F., Jackson, L.A., Kenigsberg, T.A., Kuckler, L., Tseng, H.F., Williams, J.T.B., et al. (2025). Differences in influenza vaccination coverage by race and ethnicity across age groups in the Vaccine Safety Datalink, 2017–18 through 2022–23 influenza seasons. Vaccine 64, 127667. 10.1016/j.vaccine.2025.127667.

14. Kumar, B.V., Connors, T.J., and Farber, D.L. (2018). Human T Cell Development, Localization, and Function throughout Life. Immunity 48, 202–213. 10.1016/j.immuni.2018.01.007.

15. Choi, J.O., Seo, Y., and Hwang, S.S. (2025). Guardians of silence: transcriptional networks in T cell quiescence. Exp Mol Med 57, 1663–1672. 10.1038/s12276-025-01516-y.

16. Lumnitzer, M.E., Lovell, S.A., Condotta, S.A., and Richer, M.J. (2025). Restraining the killers: regulation of T cell quiescence. Trends in Immunology 46, 525–535. 10.1016/j.it.2025.04.008.

17. Adu-Berchie, K., Obuseh, F.O., and Mooney, D.J. (2023). T Cell Development and Function. Rejuvenation Research 26, 126–138. 10.1089/rej.2023.0015.

18. Kaech, S.M., and Cui, W. (2012). Transcriptional control of effector and memory CD8+ T cell differentiation. Nat Rev Immunol 12, 749–761. 10.1038/nri3307.

19. Siu, J.H.Y., Coelho, S., Palomeras, A., Belij-Rammerstorfer, S., Barman, N., Lee, C.H., Ströbel, T., Thorpe, C.J., Kaur, C., Cole, T., et al. (2025). Early lymph node T follicular helper cell signalling hub drives influenza vaccine response in an ancestrally diverse cohort. eBioMedicine 122. 10.1016/j.ebiom.2025.106036.

20. Auton, A., Abecasis, G.R., Altshuler, D.M., Durbin, R.M., Abecasis, G.R., Bentley, D.R., Chakravarti, A., Clark, A.G., Donnelly, P., Eichler, E.E., et al. (2015). A global reference for human genetic variation. Nature 526, 68–74. 10.1038/nature15393.

21. Bergström, A., McCarthy, S.A., Hui, R., Almarri, M.A., Ayub, Q., Danecek, P., Chen, Y., Felkel, S., Hallast, P., Kamm, J., et al. (2020). Insights into human genetic variation and population history from 929 diverse genomes. Science 367, eaay5012. 10.1126/science.aay5012.

22. Alexander, D.H., Novembre, J., and Lange, K. (2009). Fast model-based estimation of ancestry in unrelated individuals. Genome Res. 19, 1655–1664. 10.1101/gr.094052.109.

23. Gonzalez-Galarza, F.F., McCabe, A., Santos, E.J.M.D., Jones, J., Takeshita, L., Ortega-Rivera, N.D., Cid-Pavon, G.M.D., Ramsbottom, K., Ghattaoraya, G., Alfirevic, A., et al. (2019). Allele frequency net database (AFND) 2020 update: gold-standard data classification, open access genotype data and new query tools. Nucleic Acids Research, gkz1029. 10.1093/nar/gkz1029.

24. Newton, R.H., Shrestha, S., Sullivan, J.M., Yates, K.B., Compeer, E.B., Ron-Harel, N., Blazar, B.R., Bensinger, S.J., Haining, W.N., Dustin, M.L., et al. (2018). Maintenance of CD4 T cell fitness through regulation of Foxo1. Nat Immunol 19, 838–848. 10.1038/s41590-018-0157-4.

25. Roychoudhuri, R., Clever, D., Li, P., Wakabayashi, Y., Quinn, K.M., Klebanoff, C.A., Ji, Y., Sukumar, M., Eil, R.L., Yu, Z., et al. (2016). BACH2 regulates CD8(+) T cell differentiation by controlling access of AP-1 factors to enhancers. Nat Immunol 17, 851–860. 10.1038/ni.3441.

26. Yao, C., Lou, G., Sun, H.-W., Zhu, Z., Sun, Y., Chen, Z., Chauss, D., Moseman, E.A., Cheng, J., D’Antonio, M.A., et al. (2021). BACH2 enforces the transcriptional and epigenetic programs of stem-like CD8+ T cells. Nat Immunol 22, 370–380. 10.1038/s41590-021-00868-7.

27. Rebuffet, L., Melsen, J.E., Escalière, B., Basurto-Lozada, D., Bhandoola, A., Björkström, N.K., Bryceson, Y.T., Castriconi, R., Cichocki, F., Colonna, M., et al. (2024). High-dimensional single-cell analysis of human natural killer cell heterogeneity. Nat Immunol 25, 1474–1488. 10.1038/s41590-024-01883-0.

28. Schattgen, S.A., Turner, J.S., Ghonim, M.A., Crawford, J.C., Schmitz, A.J., Kim, H., Zhou, J.Q., Awad, W., Mettelman, R.C., Kim, W., et al. (2024). Influenza vaccination stimulates maturation of the human T follicular helper cell response. Nat Immunol 25, 1742–1753. 10.1038/s41590-024-01926-6.

29. Joshi-Tope, G., Gillespie, M., Vastrik, I., D’Eustachio, P., Schmidt, E., de Bono, B., Jassal, B., Gopinath, G.R., Wu, G.R., Matthews, L., et al. (2005). Reactome: a knowledgebase of biological pathways. Nucleic Acids Res 33, D428–D432. 10.1093/nar/gki072.

30. Liang, H., Tang, J., Liu, Z., Liu, Y., Huang, Y., Xu, Y., Hao, P., Yin, Z., Zhong, J., Ye, L., et al. (2019). ZIKV infection induces robust Th1-like Tfh cell and long-term protective antibody responses in immunocompetent mice. Nat Commun 10, 3859. 10.1038/s41467-019-11754-0.

31. Horiuchi, S., Wu, H., Liu, W.-C., Schmitt, N., Provot, J., Liu, Y., Bentebibel, S.-E., Albrecht, R.A., Schotsaert, M., Forst, C.V., et al. (2021). Tox2 is required for the maintenance of GC T_FH_ cells and the generation of memory T_FH_ cells. Sci. Adv. 7, eabj1249. 10.1126/sciadv.abj1249.

32. Mitra, M., Batista, S.L., and Coller, H.A. (2025). Transcription factor networks in cellular quiescence. Nat Cell Biol 27, 14–27. 10.1038/s41556-024-01582-w.

33. Azevedo, C.M., Xie, B., Gunn, W.G., Peralta, R.M., Dantas, C.S., Fernandes-Mendes, H., Joshi, S., Dean, V., Almeida, P., Wilfahrt, D., et al. (2025). Reprogramming CD8+ T-cell Branched *N-* Glycosylation Limits Exhaustion, Enhancing Cytotoxicity and Tumor Killing. Cancer Immunology Research 13, 1655–1673. 10.1158/2326-6066.CIR-25-0313.

34. Cheng, J., Xiao, Y., Peng, T., Zhang, Z., Qin, Y., Wang, Y., Shi, J., Yan, J., Zhao, Z., Zheng, L., et al. (2025). ETV7 limits the antiviral and antitumor efficacy of CD8+ T cells by diverting their fate toward exhaustion. Nat Cancer 6, 338–356. 10.1038/s43018-024-00892-0.

35. Basmaciogullari, S., and Pizzato, M. (2014). The activity of Nef on HIV-1 infectivity. Front. Microbiol. 5. 10.3389/fmicb.2014.00232.

36. Kotliar, D., Curtis, M., Agnew, R., Weinand, K., Nathan, A., Baglaenko, Y., Slowikowski, K., Zhao, Y., Sabeti, P.C., Rao, D.A., et al. (2025). Reproducible single-cell annotation of programs underlying T cell subsets, activation states and functions. Nat Methods 22, 1964–1980. 10.1038/s41592-025-02793-1.

37. Chu, Y., Dai, E., Li, Y., Han, G., Pei, G., Ingram, D.R., Thakkar, K., Qin, J.-J., Dang, M., Le, X., et al. (2023). Pan-cancer T cell atlas links a cellular stress response state to immunotherapy resistance. Nat Med 29, 1550–1562. 10.1038/s41591-023-02371-y.

38. Vogler, M., Braun, Y., Smith, V.M., Westhoff, M.-A., Pereira, R.S., Pieper, N.M., Anders, M., Callens, M., Vervliet, T., Abbas, M., et al. (2025). The BCL2 family: from apoptosis mechanisms to new advances in targeted therapy. Sig Transduct Target Ther 10, 91. 10.1038/s41392-025-02176-0.

39. Faure, L., Soldatov, R., Kharchenko, P.V., and Adameyko, I. (2023). scFates: a scalable python package for advanced pseudotime and bifurcation analysis from single-cell data. Bioinformatics 39, btac746. 10.1093/bioinformatics/btac746.

40. Galluzzi, L. (2025). T cell exhaustion: early or late in tumour progression? Nat Rev Immunol 25, 227–228. 10.1038/s41577-025-01158-1.

41. Setty, M., Kiseliovas, V., Levine, J., Gayoso, A., Mazutis, L., and Pe’er, D. (2019). Characterization of cell fate probabilities in single-cell data with Palantir. Nat Biotechnol 37, 451– 460. 10.1038/s41587-019-0068-4.

42. Van Houcke, M., Wuyts, S., Bosschaerts, T., Chiffelle, J., Auger, A., Coukos, G., Harari, A., and Meysman, P. (2026). Applications of T-cell receptor specificity annotation models for quality control and immunomonitoring in adoptive T-cell therapies. ImmunoInformatics 22, 100067. 10.1016/j.immuno.2026.100067.

43. Mayer-Blackwell, K., Schattgen, S., Cohen-Lavi, L., Crawford, J.C., Souquette, A., Gaevert, J.A., Hertz, T., Thomas, P.G., Bradley, P., and Fiore-Gartland, A. (2021). TCR meta-clonotypes for biomarker discovery with tcrdist3 enabled identification of public, HLA-restricted clusters of SARS-CoV-2 TCRs. eLife 10, e68605. 10.7554/eLife.68605.

44. Kleshchevnikov, V., Shmatko, A., Dann, E., Aivazidis, A., King, H.W., Li, T., Elmentaite, R., Lomakin, A., Kedlian, V., Gayoso, A., et al. (2022). Cell2location maps fine-grained cell types in spatial transcriptomics. Nat Biotechnol 40, 661–671. 10.1038/s41587-021-01139-4.

45. Nakaya, H.I., Hagan, T., Duraisingham, S.S., Lee, E.K., Kwissa, M., Rouphael, N., Frasca, D., Gersten, M., Mehta, A.K., Gaujoux, R., et al. (2015). Systems Analysis of Immunity to Influenza Vaccination across Multiple Years and in Diverse Populations Reveals Shared Molecular Signatures. Immunity 43, 1186–1198. 10.1016/j.immuni.2015.11.012.

46. Tsang, J.S., Schwartzberg, P.L., Kotliarov, Y., Biancotto, A., Xie, Z., Germain, R.N., Wang, E., Olnes, M.J., Narayanan, M., Golding, H., et al. (2014). Global Analyses of Human Immune Variation Reveal Baseline Predictors of Postvaccination Responses. Cell 157, 499–513. 10.1016/j.cell.2014.03.031.

47. Brodin, P., Jojic, V., Gao, T., Bhattacharya, S., Angel, C.J.L., Furman, D., Shen-Orr, S., Dekker, C.L., Swan, G.E., Butte, A.J., et al. (2015). Variation in the Human Immune System Is Largely Driven by Non-Heritable Influences. Cell 160, 37–47. 10.1016/j.cell.2014.12.020.

48. Corcoran, M., Narang, S., Kaduk, M., Chernyshev, M., Färnert, A., Sundling, C., and Karlsson Hedestam, G.B. (2026). Ultra-high-throughput IGH genotyping of 25 global populations reveals population-biased allelic diversity and homozygous V and D gene deletions. Immunity 59, 1107–1122.e5. 10.1016/j.immuni.2026.01.026.

49. Law, H., Mach, M., Howe, A., Obeid, S., Milner, B., Carey, C., Elfis, M., Fsadni, B., Ognenovska, K., Phan, T.G., et al. (2022). Early expansion of CD38+ICOS+ GC Tfh in draining lymph nodes during influenza vaccination immune response. iScience 25. 10.1016/j.isci.2021.103656.

50. Zoldan, K., Ehrlich, S., Killmer, S., Wild, K., Smits, M., Russ, M., Globig, A.-M., Hofmann, M., Thimme, R., and Boettler, T. (2021). Th1-Biased Hepatitis C Virus-Specific Follicular T Helper-Like Cells Effectively Support B Cells After Antiviral Therapy. Front. Immunol. 12, 742061. 10.3389/fimmu.2021.742061.

51. Bentebibel, S.-E., Khurana, S., Schmitt, N., Kurup, P., Mueller, C., Obermoser, G., Palucka, A.K., Albrecht, R.A., Garcia-Sastre, A., Golding, H., et al. (2016). ICOS+PD-1+CXCR3+ T follicular helper cells contribute to the generation of high-avidity antibodies following influenza vaccination. Sci Rep 6, 26494. 10.1038/srep26494.

52. Bentebibel, S.-E., Lopez, S., Obermoser, G., Schmitt, N., Mueller, C., Harrod, C., Flano, E., Mejias, A., Albrecht, R.A., Blankenship, D., et al. (2013). Induction of ICOS^+^ CXCR3^+^ CXCR5^+^ T_H_ Cells Correlates with Antibody Responses to Influenza Vaccination. Sci. Transl. Med. 5. 10.1126/scitranslmed.3005191.

53. Bosma, D.M.T., Busselaar, J., Staal, M.D., Koning, M. de, Reljić, M., Lei, X., Wit, T. de, Xiao, Y., Borst, J., and Salerno, F. (2026). Requirements for development of T helper 1 and T follicular helper cells from a common precursor. Cell Reports 45. 10.1016/j.celrep.2026.117296.

54. Locci, M., Havenar-Daughton, C., Landais, E., Wu, J., Kroenke, M.A., Arlehamn, C.L., Su, L.F., Cubas, R., Davis, M.M., Sette, A., et al. (2013). Human Circulating PD-1+CXCR3−CXCR5+ Memory Tfh Cells Are Highly Functional and Correlate with Broadly Neutralizing HIV Antibody Responses. Immunity 39, 758–769. 10.1016/j.immuni.2013.08.031.

55. He, R., Zheng, X., Zhang, J., Liu, B., Wang, Q., Wu, Q., Liu, Z., Chang, F., Hu, Y., Xie, T., et al. (2023). SARS-CoV-2 spike-specific TFH cells exhibit unique responses in infected and vaccinated individuals. Sig Transduct Target Ther 8, 393. 10.1038/s41392-023-01650-x.

56. Hwang, S.S., Lim, J., Yu, Z., Kong, P., Sefik, E., Xu, H., Harman, C.C.D., Kim, L.K., Lee, G.R., Li, H.-B., et al. (2020). mRNA destabilization by BTG1 and BTG2 maintains T cell quiescence. Science 367, 1255–1260. 10.1126/science.aax0194.

57. Sidwell, T., and Kallies, A. (2016). Bach2 is required for B cell and T cell memory differentiation. Nat Immunol 17, 744–745. 10.1038/ni.3493.

58. Hu, T., Zhu, Z., Luo, Y., Wizzard, S., Hoar, J., Shinde, S.S., Yihunie, K., Yao, C., and Wu, T. (2026). BACH2 dosage establishes the hierarchy of stemness and fine-tunes antitumor immunity in CAR T cells. Nat Immunol 27, 425–435. 10.1038/s41590-025-02388-0.

59. Conti, A.G., Evans, A.C., von Linde, T., Deguit, C.D.T., Whiteside, S.K., Wesolowski, A.J., Imianowski, C.J., Yamashita-Kanemaru, Y., Dahmani, L., Chapman, J., et al. (2026). Fine-tuning BACH2 dosage balances stemness and effector function to enhance antitumor T cell therapy. Nat Immunol 27, 436–451. 10.1038/s41590-025-02389-z.

60. Gebhardt, T., Park, S.L., and Parish, I.A. (2023). Stem-like exhausted and memory CD8+ T cells in cancer. Nat Rev Cancer 23, 780–798. 10.1038/s41568-023-00615-0.

61. Wei, Y., Ma, H.K., Wong, M.E., Back, H., Papasavvas, E., Mounzer, K., Aberra, F., Morgenstern, R., Tebas, P., Konnikova, L., et al. (2025). Transcription factor BACH2 shapes tissue-resident memory T cell programs to promote HIV-1 persistence. Immunity 58, 2878–2898.e11. 10.1016/j.immuni.2025.07.022.

62. Karin, N. (2020). CXCR3 Ligands in Cancer and Autoimmunity, Chemoattraction of Effector T Cells, and Beyond. Front. Immunol. 11, 976. 10.3389/fimmu.2020.00976.

63. Groom, J.R., Richmond, J., Murooka, T.T., Sorensen, E.W., Sung, J.H., Bankert, K., von Andrian, U.H., Moon, J.J., Mempel, T.R., and Luster, A.D. (2012). CXCR3 Chemokine Receptor-Ligand Interactions in the Lymph Node Optimize CD4+ T Helper 1 Cell Differentiation. Immunity 37, 1091– 1103. 10.1016/j.immuni.2012.08.016.

64. Gerlach, C., Moseman, E.A., Loughhead, S.M., Alvarez, D., Zwijnenburg, A.J., Waanders, L., Garg, R., Torre, J.C. de la, and Andrian, U.H. von (2016). The Chemokine Receptor CX3CR1 Defines Three Antigen-Experienced CD8 T Cell Subsets with Distinct Roles in Immune Surveillance and Homeostasis. Immunity 45, 1270–1284. 10.1016/j.immuni.2016.10.018.

65. Braun, A., Worbs, T., Moschovakis, G.L., Halle, S., Hoffmann, K., Bölter, J., Münk, A., and Förster, R. (2011). Afferent lymph–derived T cells and DCs use different chemokine receptor CCR7– dependent routes for entry into the lymph node and intranodal migration. Nat Immunol 12, 879– 887. 10.1038/ni.2085.

66. Allenspach, E.J., Lemos, M.P., Porrett, P.M., Turka, L.A., and Laufer, T.M. (2008). Migratory and Lymphoid-Resident Dendritic Cells Cooperate to Efficiently Prime Naive CD4 T cells. Immunity 29, 795–806. 10.1016/j.immuni.2008.08.013.

67. Zohora, F.T., Paliwal, D., Flores-Figueroa, E., Li, J., Gao, T., Notta, F., and Schwartz, G.W. (2025). CellNEST reveals cell–cell relay networks using attention mechanisms on spatial transcriptomics. Nat Methods 22, 1505–1519. 10.1038/s41592-025-02721-3.

68. Yoon, I.-S., Park, H., Kwak, H.-W., Woo Jung, Y., and Nam, J.-H. (2017). Macrophage-derived insulin-like growth factor-1 affects influenza vaccine efficacy through the regulation of immune cell homeostasis. Vaccine 35, 4687–4694. 10.1016/j.vaccine.2017.07.037.

69. Chang, C.C., Chow, C.C., Tellier, L.C., Vattikuti, S., Purcell, S.M., and Lee, J.J. (2015). Second-generation PLINK: rising to the challenge of larger and richer datasets. Gigascience 4, s13742–015-0047–0048. 10.1186/s13742-015-0047-8.

70. Purcell, S., Neale, B., Todd-Brown, K., Thomas, L., Ferreira, M.A.R., Bender, D., Maller, J., Sklar, P., Bakker, P.I.W. de, Daly, M.J., et al. (2007). PLINK: A Tool Set for Whole-Genome Association and Population-Based Linkage Analyses. The American Journal of Human Genetics 81, 559–575. 10.1086/519795.

71. Ziyatdinov, A., Torres, J., Alegre-Díaz, J., Backman, J., Mbatchou, J., Turner, M., Gaynor, S.M., Joseph, T., Zou, Y., Liu, D., et al. (2023). Genotyping, sequencing and analysis of 140,000 adults from Mexico City. Nature 622, 784–793. 10.1038/s41586-023-06595-3.

72. Bernstein, N.J., Fong, N.L., Lam, I., Roy, M.A., Hendrickson, D.G., and Kelley, D.R. (2020). Solo: Doublet Identification in Single-Cell RNA-Seq via Semi-Supervised Deep Learning. cels 11, 95–101.e5. 10.1016/j.cels.2020.05.010.

73. Heaton, H., Talman, A.M., Knights, A., Imaz, M., Gaffney, D.J., Durbin, R., Hemberg, M., and Lawniczak, M.K.N. (2020). Souporcell: robust clustering of single-cell RNA-seq data by genotype without reference genotypes. Nat Methods 17, 615–620. 10.1038/s41592-020-0820-1.

74. Wolf, F.A., Angerer, P., and Theis, F.J. (2018). SCANPY: large-scale single-cell gene expression data analysis. Genome Biol 19, 15. 10.1186/s13059-017-1382-0.

75. Wolock, S.L., Lopez, R., and Klein, A.M. (2019). Scrublet: Computational Identification of Cell Doublets in Single-Cell Transcriptomic Data. Cell Syst 8, 281–291.e9. 10.1016/j.cels.2018.11.005.

76. Popescu, D.-M., Botting, R.A., Stephenson, E., Green, K., Webb, S., Jardine, L., Calderbank, E.F., Polanski, K., Goh, I., Efremova, M., et al. (2019). Decoding human fetal liver haematopoiesis. Nature 574, 365–371. 10.1038/s41586-019-1652-y.

77. Xu, C., Prete, M., Webb, S., Jardine, L., Stewart, B.J., Hoo, R., He, P., Meyer, K.B., and Teichmann, S.A. (2023). Automatic cell-type harmonization and integration across Human Cell Atlas datasets. Cell 186, 5876–5891.e20. 10.1016/j.cell.2023.11.026.

78. Boyeau, P., Hong, J., Gayoso, A., Kim, M., McFaline-Figueroa, J.L., Jordan, M.I., Azizi, E., Ergen, C., and Yosef, N. (2025). Deep generative modeling of sample-level heterogeneity in single-cell genomics. Nat Methods 22, 2264–2274. 10.1038/s41592-025-02808-x.

79. Korsunsky, I., Millard, N., Fan, J., Slowikowski, K., Zhang, F., Wei, K., Baglaenko, Y., Brenner, M., Loh, P., and Raychaudhuri, S. (2019). Fast, sensitive and accurate integration of single-cell data with Harmony. Nat Methods 16, 1289–1296. 10.1038/s41592-019-0619-0.

80. Hoffman, G.E., and Schadt, E.E. (2016). variancePartition: interpreting drivers of variation in complex gene expression studies. BMC Bioinformatics 17, 483. 10.1186/s12859-016-1323-z.

81. Badia-i-Mompel, P., Vélez Santiago, J., Braunger, J., Geiss, C., Dimitrov, D., Müller-Dott, S., Taus, P., Dugourd, A., Holland, C.H., Ramirez Flores, R.O., et al. (2022). decoupleR: ensemble of computational methods to infer biological activities from omics data. Bioinformatics Advances 2, vbac016. 10.1093/bioadv/vbac016.

82. Love, M.I., Huber, W., and Anders, S. (2014). Moderated estimation of fold change and dispersion for RNA-seq data with DESeq2. Genome Biol 15, 550. 10.1186/s13059-014-0550-8.

83. Heumos, L., Ji, Y., May, L., Green, T.D., Peidli, S., Zhang, X., Wu, X., Ostner, J., Schumacher, A., Hrovatin, K., et al. (2026). Pertpy: an end-to-end framework for perturbation analysis. Nat Methods 23, 350–359. 10.1038/s41592-025-02909-7.

84. Bürkner, P.-C. (2017). **brms** : An *R* Package for Bayesian Multilevel Models Using *Stan*. J. Stat. Soft. 80. 10.18637/jss.v080.i01.

85. Carpenter, B., Gelman, A., Hoffman, M.D., Lee, D., Goodrich, B., Betancourt, M., Brubaker, M., Guo, J., Li, P., and Riddell, A. (2017). *Stan* : A Probabilistic Programming Language. J. Stat. Soft. 76. 10.18637/jss.v076.i01.

86. Chen, Y., Chen, L., Lun, A.T.L., Baldoni, P.L., and Smyth, G.K. (2025). edgeR v4: powerful differential analysis of sequencing data with expanded functionality and improved support for small counts and larger datasets. Nucleic Acids Research 53, gkaf018. 10.1093/nar/gkaf018.

87. Fang, Z., Liu, X., and Peltz, G. (2023). GSEApy: a comprehensive package for performing gene set enrichment analysis in Python. Bioinformatics 39, btac757. 10.1093/bioinformatics/btac757.

88. Jin, S., Plikus, M.V., and Nie, Q. (2025). CellChat for systematic analysis of cell–cell communication from single-cell transcriptomics. Nat Protoc 20, 180–219. 10.1038/s41596-024-01045-4.

89. Sturm, G., Szabo, T., Fotakis, G., Haider, M., Rieder, D., Trajanoski, Z., and Finotello, F. (2020). Scirpy: a Scanpy extension for analyzing single-cell T-cell receptor-sequencing data. Bioinformatics 36, 4817–4818. 10.1093/bioinformatics/btaa611.

90. Peng, K., Moore, J., Vahed, M., Brito, J., Kao, G., Burkhardt, A.M., Alachkar, H., and Mangul, S. (2022). pyTCR: A comprehensive and scalable solution for TCR-Seq data analysis to facilitate reproducibility and rigor of immunogenomics research. Front. Immunol. 13, 954078. 10.3389/fimmu.2022.954078.

91. Arunkumar, M., and Zielinski, C.E. (2021). T-Cell Receptor Repertoire Analysis with Computational Tools—An Immunologist’s Perspective. Cells 10, 3582. 10.3390/cells10123582.

